# Quantification of the pace of biological aging in humans through a blood test: The DunedinPoAm DNA methylation algorithm

**DOI:** 10.1101/2020.02.05.927434

**Authors:** DW Belsky, A Caspi, L Arseneault, A Baccarelli, D Corcoran, X Gao, E Hannon, HL Harrington, LJH Rasmussen, R Houts, K Huffman, WE Kraus, D Kwon, J Mill, CF Pieper, J Prinz, R Poulton, J Schwartz, K Sugden, P Vokonas, BS Williams, TE Moffitt

## Abstract

Biological aging is the gradual, progressive decline in system integrity that occurs with advancing chronological age, causing morbidity and disability. Measurements of the pace of aging are needed to serve as surrogate endpoints in trials of therapies designed to prevent disease by slowing biological aging. We report a blood DNA-methylation measure that is sensitive to variation in the pace of biological aging among individuals born the same year. We first modeled longitudinal change in 18 biomarkers tracking organ-system integrity across 12 years of follow-up in the Dunedin birth cohort. Rates of change in each biomarker were composited to form a measure of aging-related decline, termed Pace of Aging. Elastic-net regression was used to develop a DNA-methylation predictor of Pace of Aging, called DunedinPoAm for Dunedin (P)ace (o)f (A)ging (m)ethylation. Validation analyses showed DunedinPoAm was associated with functional decline in the Dunedin Study and more advanced biological age in the Understanding Society Study, predicted chronic disease and mortality in the Normative Aging Study, was accelerated by early-life adversity in the E-risk Study, and DunedinPoAm prediction was disrupted by caloric restriction in the CALERIE trial. DunedinPoAm generally outperformed epigenetic clocks. Findings provide proof-of-principle for DunedinPoAm as a single-time-point measure of a person’s pace of biological aging.

## INTRODUCTION

Aging of the global population is producing forecasts of rising burden of disease and disability (Harper, 2014). Because this burden arises from multiple age-related diseases, treatments for single diseases will not address the burden challenge (Goldman et al., 2013). Geroscience research suggests an appealing alternative: treatments to slow aging itself could prevent or delay the multiple diseases that increase with advancing age, perhaps with a single therapeutic approach (Gladyshev, 2016; Kaeberlein, 2013). Aging can be understood as a gradual and progressive deterioration in biological system integrity (Kirkwood, 2005). This deterioration is thought to arise from an accumulation of cellular-level changes. These changes, in turn, increase vulnerability to diseases affecting many different organ systems (Kennedy et al., 2014; López-Otín et al., 2013). Animal studies suggest treatments that slow the accumulation of cellular-level changes can extend healthy lifespan (Campisi et al., 2019; Kaeberlein et al., 2015). However, human trials of these treatments are challenging because humans live much longer than model animals, making it time-consuming and costly to follow up human trial participants to test treatment effects on healthy lifespan. This challenge will be exacerbated in trials that will give treatments to young or middle-aged adults, with the aim to prevent the decline in system integrity that antedates disease onset by years. Involving young and midlife adults in healthspan-extension trials has been approved for development by the National Advisory Council on Aging (2019 CTAP report to NACA). In midlife trials of treatments to slow aging, called geroprotectors (Moskalev et al., 2016), traditional endpoints such as disease diagnosis or death are too far in the future to serve as outcomes. Translation of geroprotector treatments to humans could be aided by measures that quantify the pace of deterioration in biological system integrity in human aging. Such measures could be used as surrogate endpoints for healthy lifespan extension (Justice et al., 2016, 2018; Moskalev et al., 2016), even with young-to-midlife adult trial participants. A useful measure should be non-invasive, inexpensive, reliable, and highly sensitive to biological change.

Recent efforts to develop such measures have focused on blood DNA methylation as a biological substrate highly sensitive to changes in chronological age (Fahy et al., 2019; Horvath and Raj, 2018). Methylation-clock algorithms have been developed to identify methylation patterns that characterize individuals of different chronological ages. However, a limitation is that individuals born in different years have grown up under different historical conditions (Schaie, 1967). For example, people born 70 years ago experienced more exposure to childhood diseases, tobacco smoke, airborne lead, and less exposure to antibiotics and other medications, and lower quality nutrition, all of which leave signatures on DNA methylation (Bell et al., 2019). As a result, the clocks confound methylation patterns arising from early-life exposures to methylation-altering factors with methylation patterns related to biological aging during adulthood. An alternative approach is to study individuals who were all born the same year, and find methylation patterns that differentiate those who have been aging biologically faster or slower than their same-age peers. The current article reports four steps in our work toward developing a blood DNA methylation measure to represent individual variation in the pace of biological aging.

In Step 1, which we previously reported (Belsky et al., 2015), we collected a panel of 18 blood chemistry and organ-system function biomarkers at three successive waves of the Dunedin Longitudinal Study of a 1972-73 population-representative one-year birth cohort (N=1037). We used repeated-measures data collected when Study members were aged 26, 32, and 38 years old to quantify rates of biological change. We modelled the rate of change in each biomarker and calculated how each Study member’s personal rate-of-change on that biomarker differed from the cohort norm. We then combined the 18 personal rates of change across the panel of biomarkers to compute a composite for each Study member that we called the Pace of Aging. Pace of Aging represents a personal rate of multi-organ-system decline over a dozen years. Pace of Aging was normally distributed, and showed marked variation among Study members who were all the same chronological age, confirming that individual differences in biological aging do emerge already by age 38, years before chronic disease onset.

In Step 2, which we previously reported, we validated the Pace of Aging against known criteria. As compared to other Study members who were the same chronological age but had slower Pace of Aging, Study members with faster Pace of Aging performed more poorly on tests of physical function; showed signs of cognitive decline on a panel of dementia-relevant neuropsychological tests from an early-life baseline; were rated as looking older based on facial photographs; and reported themselves to be in worse health (Belsky et al., 2015). Subsequently, we reported that faster Pace of Aging is associated with early-life factors important for aging: familial longevity, low childhood social class, and adverse childhood experiences (Belsky et al., 2017a), and that faster Pace of Aging is associated with older scores on Brain Age, a machine-learning-derived measure of structural MRI differences characteristic of different age groups (Elliott et al., 2019). Notably, Pace of Aging was not well-correlated with published epigenetic age clocks, which were designed to measure how old a person is rather than how fast they are aging biologically (Belsky et al., 2018).

In Step 3, which we report here, we distill the Pace of Aging into a measurement that can be obtained from a single blood sample. Here we focused on blood DNA methylation as an accessible molecular measurement that is sensitive to changes in physiology occurring in multiple organ systems (Birney et al., 2016; Bolund et al., 2017; Chambers et al., 2015; Chu et al., 2017; Hedman Åsa K. et al., 2017; Ma et al., 2019; Mill and Heijmans, 2013; Morris et al., 2017; Wahl et al., 2017). We used data about the Pace of Aging from age 26 to 38 years in the Dunedin Study along with whole-genome methylation data at age 38 years. Elastic-net regression was applied to derive an algorithm that captured DNA methylation patterns linked with variation among individuals in their Pace of Aging. The algorithm is hereafter termed “DunedinPoAm”.

DunedinPoAm is qualitatively different from previously published DNA methylation measures of aging that were developed by comparing older individuals to younger ones. Those measures, often referred to as “clocks,” are state measures. They estimate how much aging has occurred in an individual up to the point of measurement. DunedinPoAm is a rate measure. It is based on comparison of longitudinal change over time in 18 biomarkers of organ-system integrity among individuals who are all the same chronological age. DunedinPoAm estimates how fast aging is occurring during the years leading up to the time of measurement. Rather than a clock that records how much time has passed, DunedinPoAm is designed to function as a speedometer, recording how fast the subject is aging. In Step 4, which we report here, we validated the DunedinPoAm in 5 ways. First, using the Dunedin Study, we tested if Study member’s DunedinPoAm measured when they were aged 38 years could predict deficits in physical and cognitive functioning seven years later, when the cohort was aged 45 years. Second, we applied the DunedinPoAm algorithm to DNA methylation data from a second, cross-sectional, study of adults to evaluate patterning of DunedinPoAm by chronological age and sex and to test correlations of DunedinPoAm with self-reported health and proposed measures of biological age, including three epigenetic clocks. Third, we applied the DunedinPoAm algorithm to DNA methylation data from a third, longitudinal study of older men to test associations with chronic-disease morbidity and mortality. Fourth, we applied the DunedinPoAm algorithm to DNA methylation data from a fourth, longitudinal, study of young people to test if DunedinPoAm was accelerated by exposure to poverty and victimization, factors which are known to shorten healthy lifespan. Finally, to ascertain the potential usefulness of DunedinPoAm as a measure for trials of geroprotector treatments, we applied the algorithm to DNA methylation data from a randomized trial of caloric restriction, CALERIE (Ravussin et al., 2015). Earlier we reported from this trial that the intervention (two years of prescribed 25% caloric restriction) slowed the rate of biological aging as measured by a blood-chemistry biological-age composite measure (Belsky et al., 2017b). Here, using newly generated methylation data from blood drawn at the CALERIE baseline assessment, we tested if (a) DunedinPoAm from blood drawn before caloric restriction could predict the future rate of biological aging of participants during the two-year trial, and (b) if this prediction was disrupted in participants who underwent caloric restriction, but not among control participants. We report promising results from this four-step research program, while appreciating that additional measurement development will be needed to support applied use of DunedinPoAm. A graphical illustration of our study designed is presented in **Figure 1**.

**Figure 1.**
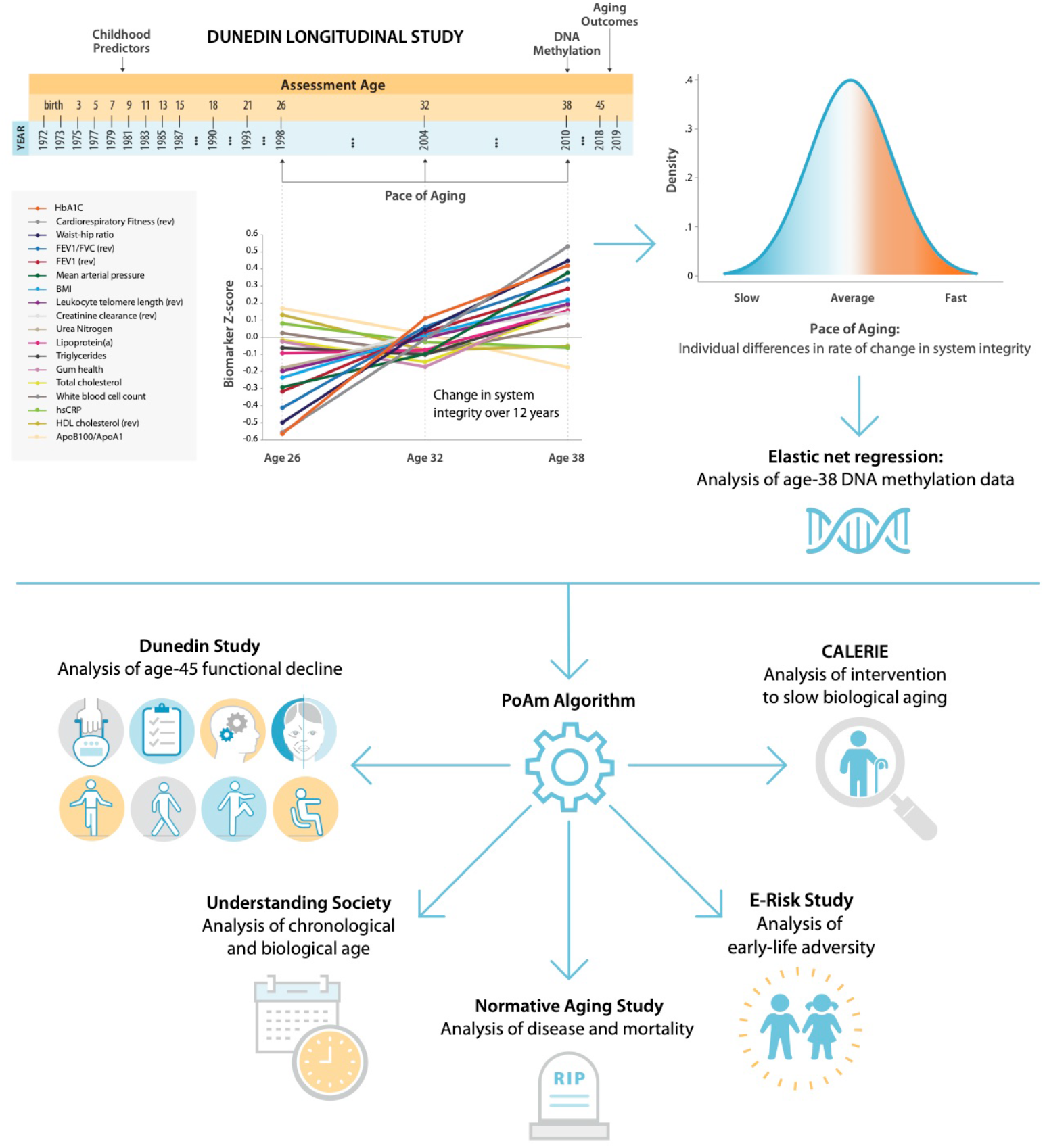
Schematic of design and follow-up of DunedinPoAm. DunedinPoAm is designed to quantify the rate of decline in system integrity experienced by an individual over the recent past; it functions like speedometer for the rate of aging. We developed DunedinPoAm from analysis of longitudinal change in 18 biomarkers of organ system integrity in the Dunedin Longitudinal Study birth cohort. Biomarkers were measured in 1998, 2004, and 2010, when all cohort members were aged 26, 32, and 38 years. We composited rates of change across the 18 biomarkers to produce a single measure of aging-related decline in system integrity, termed Pace of Aging. We then used elastic-net regression to develop a DNA-methylation predictor of Pace of Aging, called DunedinPoAm for Dunedin (P)ace (o)f (A)ging (m)ethylation. DNA methylation data for this analysis came from the age-38 assessment in 2010. We further evaluated the performance of DunedinPoAm using data from (a) the age-45 assessments of Dunedin Study members in 2018, (b) the Understanding Society Study, (c) the Normative Aging Study, (d) the E-risk Study, and (e) the CALERIE trial.

## METHODS

### Data Sources

Data were used from five studies: Dunedin Study, Understanding Society Study, the Normative Aging Study (NAS), Environmental Risk (E-Risk) Longitudinal Twin Study, and CALERIE Trial. The four datasets and measures analyzed within each of them are described in **Supplemental Materials** Section 1.

### DNA Methylation Data

DNA methylation was measured from Illumina 450k Arrays in the Dunedin Study, Normative Aging Study, and E-Risk Study and from Illumina EPIC 850k Arrays in the Understanding Society study and the CALERIE Trial. DNA was derived from whole blood samples in all studies. Dunedin Study blood draws were conducted at the cohort’s age-38 assessment during 2010-12. Understanding Society blood draws were conducted in 2012. Normative Aging Study blood draws were conducted during 1999-2013. E-Risk blood draws were conducted at the cohort’s age-18 assessment during 2012-13. CALERIE blood draws were conducted at the trial baseline assessment in 2007. Dunedin and CALERIE methylation assays were run by the Molecular Genomics Shared Resource at Duke Molecular Physiology Institute, Duke University (USA). Understanding Society and E-Risk assays were run by the Complex Disease Epigenetics Group at the University of Exeter Medical School (UK) (www.epigenomicslab.com). Normative Aging Study methylation assays were run by the Genome Research Core of the University of Illinois at Chicago. Processing protocols for the methylation data from all studies have been described previously (Dai et al., 2017; Hannon et al., 2018; Marzi et al., 2018; Panni Tommaso et al., 2016). (CALERIE data were processed according to the same protocols used for the Dunedin Study.)

#### Methylation Clocks

We computed the methylation clocks proposed by Horvath, Hannum, and Levine using the methylation data provided by the individual studies and published algorithms (Hannum et al., 2013; Horvath, 2013; Levine et al., 2018).

The Dunedin Pace of Aging methylation algorithm (DunedinPoAm) was developed using elastic-net regression analysis carried out in the Dunedin Study, as described in detail in the Results. The criterion variable was Pace of Aging. Development of the Pace of Aging is described in detail elsewhere (Belsky et al., 2015). Briefly, we conducted mixed-effects growth modeling of longitudinal change in 18 biomarkers measuring integrity of the cardiovascular, metabolic, renal, hepatic, pulmonary, periodontal, and immune systems (biomarkers are listed in the **Supplemental Materials** Section 1, description of the Dunedin Study data). For each biomarker, we estimated random slopes quantifying each participant’s own rate of change in that biomarker. We then composited slopes across the 18 biomarkers to calculate a participant’s Pace of Aging. Pace of Aging was scaled in units representing the mean trend in the cohort, i.e. the average physiological change occurring during one calendar year (N=954, M=1, SD=0.38). Of the N=819 Dunedin Study members with methylation data at age 38, N=810 had measured Pace of Aging (M=0.98, SD=0.09). This group formed the analysis sample to develop DunedinPoAm.

To compute DunedinPoAm in the Understanding Society, Normative Aging Study, E-Risk, and CALERIE Trial datasets, we applied the scoring algorithm estimated from elastic net regression in the Dunedin Study. CpG weights for the scoring algorithm are provided in **Supplemental Table 1**.

### Statistical Analysis

We conducted analysis of Dunedin, Understanding Society, NAS, E-Risk, and CALERIE data using regression models. We analyzed continuous outcome data using linear regression. We analyzed count outcome data using Poisson regression. We analyzed time-to-event outcome data using Cox proportional hazard regression. For analysis of repeated-measures longitudinal DNA methylation data in the Normative Aging Study, we used generalized estimating equations to account for non-independence of repeated observations of individuals (Ballinger, 2004), following the method in previous analysis of those data (Gao et al., 2018), and econometric fixed-effects regression (Wooldridge, 2012) to test within-person change over time. For analysis in E-Risk, which include data on twin siblings, we clustered standard errors at the family level to account for non-independence of data. For analysis of longitudinal change in clinical-biomarker biological age in CALERIE, we used mixed-effects growth models (Singer and Willett, 2003) following the method in our original analysis of those data (Belsky et al., 2017b). For regression analysis, methylation measures were adjusted for batch effects by regressing the measure on batch controls and predicting residual values. Dunedin Study, Understanding Society, E-Risk, and CALERIE analyses included covariate adjustment for sex (the Normative Aging Study included only men). Understanding Society, Normative Aging Study, and CALERIE analyses included covariate adjustment for chronological age. (Dunedin and E-Risk are birth-cohort studies and participants are all the same chronological age.) Sensitivity analyses testing covariate adjustment for estimated leukocyte distributions and smoking are reported in **Supplemental Tables 3-7**.

## RESULTS

### Capturing Pace of Aging in a single blood test

We derived the DunedinPoAm algorithm using data from Dunedin Study members for whom age-38 DNA methylation data were available (N=810). We applied elastic-net regression (Zou and Hastie, 2005) using Pace of Aging between ages 26 to 38 years as the criterion. We included all methylation probes that appear on both the Illumina 450k and EPIC arrays as potential predictor variables. We selected this overlapping set of probes for our analysis to facilitate application of the derived algorithm by other research teams using either chip. We fixed the alpha parameter to 0.5, following the approach reported by Horvath (Horvath, 2013). This analysis selected a set of 46 CpG sites (**Supplemental Table 1**, **Supplemental Materials** Section 2). The 46-CpG elastic-net-derived DunedinPoAm algorithm, applied in the age-38 Dunedin DNA methylation data, was associated with the longitudinal 26-38 Pace of Aging measure (Pearson r=0.56, **Supplemental Figure 1**). This is likely an overestimate of the true out-of-sample correlation because the analysis is based on the same data used to develop the DunedinPoAm algorithm; bootstrap-cross-validation analysis estimated the out-of-sample correlation to be r=0.33; **Supplemental Materials** Section 3).

### DunedinPoAm in midlife predicted future functional limitations

#### Physical Functioning

As a primary criterion validity analysis of DunedinPoAm, we tested prospective associations of Dunedin Study members’ age-38 DunedinPoAm values with their performance seven years later, when they were aged 45 years, on tests of balance, walking speed, chair stands, grip strength, motor coordination, and Study-member reports about physical limitations. Performance scores were reversed so that positive correlations indicated an association between faster DunedinPoAm and worse physical performance. Study members with faster DunedinPoAm at age 38 performed more poorly at age 45 on all physical performance tests, with the exception of grip strength, and reported more functional limitations (standardized effect-sizes for tests of balance, walking speed, chair stands, and physical limitations r=0.15-0.29, p<0.001 for all; grip strength r=0.05, 95% CI [−0.02-0.12], p=0.162). Effect-sizes are graphed in the dark blue bars in **Figure 2, Panel A**.

**Figure 2.**
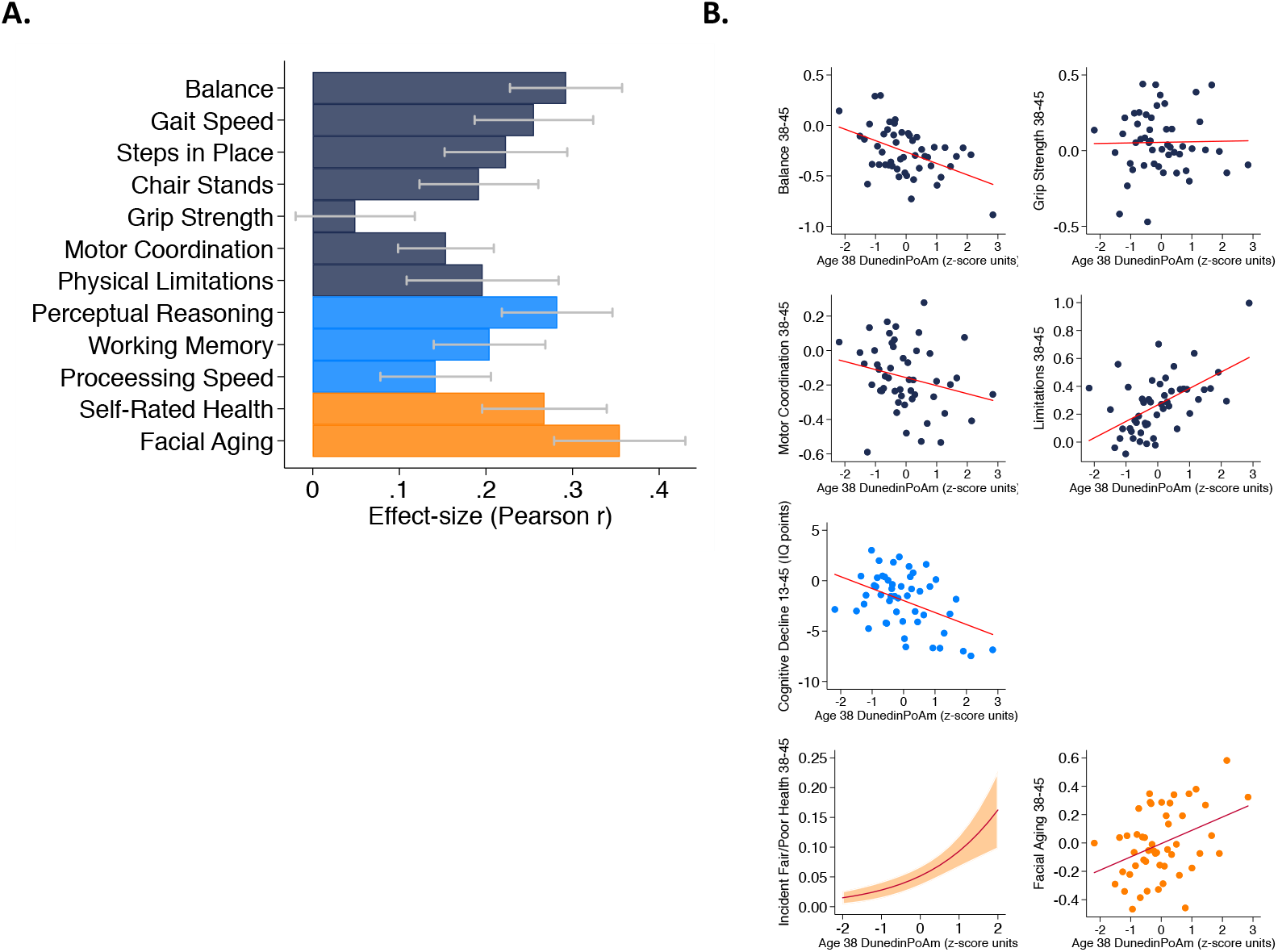
Faster age-38 DunedinPoAm is associated with poorer physical and cognitive functioning and subjective signs of aging at age 45 years, and with physical, cognitive, and subjective decline in the Dunedin Study. Panel A graphs effect-sizes for age-38 DunedinPoAm associations with age-45 measures of physical and cognitive functioning and subjective signs of aging in the Dunedin Study. Effect-sizes are standardized regression coefficients interpretable as Pearson r. Models included covariate adjustment for sex. Panel B graphs associations between DunedinPoAm and change in physical functioning between age 38 and age 45 (top row), change in cognitive functioning between age 13 and age 45 (bottom row, left-side cell), and incident fair/poor health and accelerated facial aging between ages 38 and 45 (bottom row, center and right-side cells). Graphs for changes in balance, grip-strength, physical limitations, cognition, and facial aging are binned scatterplots. Plotted points reflect average x- and y-coordinates for “bins” of approximately ten Study members. Fitted slopes show the association estimated from the raw, un-binned data. The y-axis scale on graphs of balance, grip-strength, and physical limitations shows change scores (age 45 – age 38) scaled in terms of age-38 standard deviation units. The y-axis scale on the graph of cognitive change shows the difference in IQ score (age 45 – baseline). The graph of change in facial aging shows the change in z-score between measurement intervals (age 45 – age 38). Effect-sizes reported on the graphs are standardized regression coefficients interpretable as Pearson r. Models included covariate adjustment for sex. The graph for self-rated health plots the predicted probability (fitted slope) and 95% confidence interval (shaded area) of incident fair/poor health at age 45. The effect-size reported on the graph is the incidence-rate ratio (IRR) associated with a 1-SD increase in DunedinPoAm estimated from Poisson regression. The model included covariate adjustment for sex.

#### Physical Decline

For balance, grip strength, motor coordination, and functional limitations, the Dunedin study administered the same assessments at the age-38 and age-45 assessments. We used these data to measure change in physical function across the 7-year interval. We computed change scores by subtracting the age-45 score from the age-38 score. Change scores for balance and physical limitations indicated worsening of physical functioning across the 7-year interval (in terms of age-38 standard-deviation units (SDs): balance declined by 0.26 SDs 95% CI [0.19-0.33], motor coordination declined by 0.16 [0.10-0.22] SDs, and physical limitations increased by 0.26 [0.18-0.33] SDs). In contrast, grip strength increased slightly (0.05 [0.00-0.11] SDs). Study members with faster age-38 DunedinPoAm experienced greater decline in balance at age 45 (r=0.11 [0.04-0.19], p=0.008) and a greater increase in physical limitations (r=0.10 [0.02-0.18], p=0.012). The association with change in motor coordination was consistent in direction, but was not statistically different from zero at the alpha=0.05 threshold (r=0.06 [−0.14-0.01], p=0.109). There was no association between DunedinPoAm and change in grip strength (r=0.00 [−0.07-0.08], p=0.897). Effect-sizes are graphed in the top row of **Figure 2, Panel B**.

#### Cognitive Functioning

We evaluated cognitive functioning from tests of perceptual reasoning, working memory, and processing speed, which are known to show aging related declines already by the fifth decade of life (Hartshorne and Germine, 2015; Park et al., 2002). Study members with faster age-38 DunedinPoAm performed more poorly on all age-45 cognitive tests (r=0.14-0.28, p<0.001 for all). Effect-sizes are graphed in the light-blue bars of **Figure 2, Panel A**.

#### Cognitive Decline

Cognitive functioning in early life is a potent risk factor for chronic disease and dementia in later life and for accelerated aging in midlife (Belsky et al., 2017a; Deary and Batty, 2006). Therefore, to evaluate whether associations between DunedinPoAm and cognitive test performance at age 45 might reflect reverse causation instead of early cognitive decline, we next conducted analysis of cognitive decline between adolescence and midlife. We evaluated cognitive decline by comparing Study-members’ cognitive-test performance at age 45 to their cognitive-test performance three decades earlier when they were ages 7-13 years. Cognitive performance was measured from composite scores on the Wechsler Intelligence Scales (the Wechsler Adult Intelligence Scales Version IV at the age-45 assessment and the Wechsler Intelligence Scales for Children Version R at the earlier timepoints). On average, Study members showed a decline of 2.00 IQ points (95% CI [1.31-2.70]) across the follow-up interval. We conducted two analyses to test if participants with faster DunedinPoAm experienced more cognitive decline. First, we computed difference scores (age-45 IQ – childhood baseline IQ) and regressed these difference scores on DunedinPoAm. Second, we conducted analysis of residualized change by regressing age-45 IQ on DunedinPoAm and childhood IQ. Both analyses found that Study members with faster DunedinPoAm experienced more decline (difference-score r=0.12, [0.05-0.19], p=0.001; residualized change r=0.20 [0.13-0.27]). The DunedinPoAm association with cognitive decline is graphed in the bottom-left cell of **Figure 2, Panel B**.

#### Subjective Signs of Aging

We evaluated subjective signs of aging from Study members’ ratings of their current health status (excellent, very-good, good, fair, poor) and from ratings of perceived age made by undergraduate raters based on facial photographs. Study members with faster age-38 DunedinPoAm rated themselves to be in worse health at age 45 (r=0.27, 95% CI [0.20-0.34]). These Study members were also rated as looking older (r=0.35 [0.28-0.43]). Effect-sizes are graphed in the orange bars in **Figure 2, Panel A**.

#### Subjective Signs of Decline with Aging

We next analyzed change in subjective signs of aging. Across the 7-year follow-up interval, an increasing number of Study members rated themselves as being in fair or poor health (6% rated their health as fair or poor at age 38, as compared to 8% 7 years later at age 45). Those with faster age-38 DunedinPoAm were more likely to transition to the fair/poor categories (Incidence Rate Ratio (IRR)=1.79 95% CI [1.48-2.18]). We tested if Study members with faster DunedinPoAm experienced more rapid facial aging by subtracting the age-45 score from the age 38-score and regressing this difference on DunedinPoAm. This analysis tested if Study members with faster DunedinPoAm experienced upward rank mobility within the cohort in terms of how old they looked. Study members with faster age-38 DunedinPoAm were rated as looking older relative to peers at age 45 than they had been at age 38 (r=0.10 [0.03-0.18]). Effect-sizes are graphed in the bottom-right two panels of **Figure 2, Panel B**.

#### Comparing DunedinPoAm versus Pace of Aging

We compared DunedinPoAm effect-sizes to effect-sizes for the original, 18-biomarker 3-time-point Pace of Aging. Across the domains of physical function, cognitive function, and subjective signs of aging, DunedinPoAm effect-sizes were similar to and sometimes larger than effect-sizes for the original Pace of Aging measure (**Supplemental Table 2, Supplemental Figure 1**).

Covariate adjustment to models for estimated cell counts (Houseman et al., 2012) and did not change results. Covariate adjustment for smoking history at age 38 years modestly attenuated some effect-sizes and attenuated DunedinPoAm associations with cognitive decline to near zero. Results for all models are reported in **Supplemental Table 3**.

In comparison to DunedinPoAm, effect-sizes for associations with functional limitations were smaller for the Horvath, Hannum, and Levine epigenetic clocks and, in the cases of the Horvath and Hannum clocks, were not statistically different from zero at the alpha=0.05 threshold for most outcomes. Effect-sizes are reported in **Supplemental Table 3** and plotted in **Supplemental Figure 1**.

### Evaluating DunedinPoAm and other methylation clocks in the Understanding Society study

To test variation in DunedinPoAm and to compare it with published methylation measures of biological aging, we conducted analysis using data on N=1,175 participants aged 28-95 years (M=58, SD=15; 42% male) in the UK Understanding Society cohort. In this mixed-age sample, the mean DunedinPoAm was 1.03 years of biological aging per each calendar year (SD=0.07).

We first tested if higher DunedinPoAm levels, which indicate faster aging, were correlated with older chronological age. Mortality rates increase with advancing chronological age, although there may be some slowing at the oldest ages (Barbi et al., 2018). This suggests the hypothesis that the rate of aging increases across much of the adult lifespan. Consistent with this hypothesis, Understanding Society participants who were of older chronological age tended to have faster DunedinPoAm (r=0.11, [0.06-0.17], p<0.001; **Figure 3 Panel A**). We also compared DunedinPoAm with three methylation measures of biological age: the epigenetic clocks proposed by Horvath, Hannum, and Levine (Hannum et al., 2013; Horvath, 2013; Levine et al., 2018). These epigenetic clocks were highly correlated with chronological age in the Understanding Society sample (Horvath Clock r=0.91, Hannum Clock r=0.92, Levine Clock r=0.88).

**Figure 3.**
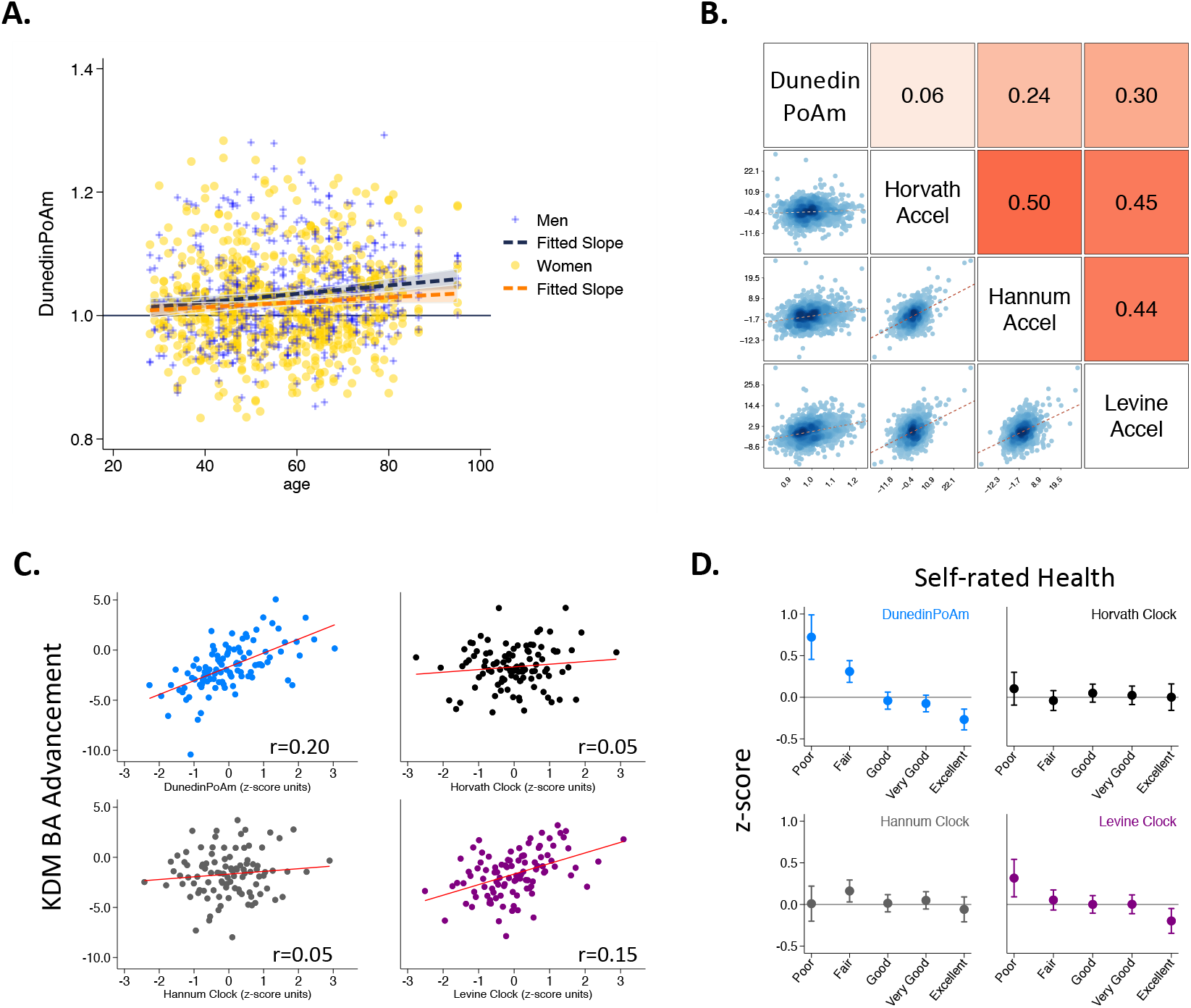
Associations among DunedinPoAm, and chronological age in the Understanding Society Study. Panel A shows a scatterplot and fitted slopes illustrating the association between chronological age (x-axis) and DunedinPoAm (y-axis) in women and men in the Understanding Society sample. Data for women are plotted with yellow dots (orange slope) and for men with blue crosses (navy slope). The figure illustrates a positive association between chronological age and DunedinPoAm (Pearson r=0.11 95% CI [0.06-0.17]). Panel B shows a matrix of association plots among DunedinPoAm and age-acceleration residuals of the Horvath, Hannum, and Levine epigenetic clocks. The diagonal cells of the matrix list the DNA methylation measures. The lower half of the matrix shows scatter plots of associations. For each scatter-plot cell, the x-axis corresponds to the variable named along the matrix diagonal to the right of the plot and the y-axis corresponds to the variable named along the matrix diagonal above the plot. The upper half of the matrix lists Pearson correlations between the DNA methylation measures. For each correlation cell, the value reflects the correlation of the variables named along the matrix diagonal to the left of the cell and below the cell. Panel C graphs binned scatterplots of associations of DunedinPoAm and epigenetic clocks with KDM Biological Age advancement (the difference between KDM Biological Age and chronological age). Each plotted point shows average x- and y-coordinates for “bins” of approximately 50 participants. Regression slopes are graphed from the raw, un-binned data. Panel D plots average values of the DNA methylation variables by Understanding Society participants’ self-rated health status. Error bars show 95% confidence intervals.

Next, to test if DunedinPoAm captured similar information about aging to published epigenetic clocks, we regressed each of the published clocks on chronological age and predicted residual values, following the procedure used by the developers of the clocks. These residuals are referred to in the literature as measures of “epigenetic age acceleration.” None of the 46 CpGs included in the DunedinPoAm algorithm overlapped with CpGs in these epigenetic clocks. Nevertheless, DunedinPoAm was moderately correlated with epigenetic age acceleration measured from the clocks proposed by Hannum (r=0.24) and Levine (r=0.30). DunedinPoAm was less-well correlated with acceleration measured from the Horvath clock (r=0.06). Associations among DunedinPoAm and the epigenetic clocks in the Understanding Society sample are shown in **Figure 3 Panel B.**

Finally, we tested correlations of DunedinPoAm with (a) a measure of biological age derived from blood chemistry and blood pressure data, and (b) a measure of self-rated health. We computed biological age from Understanding Society blood chemistry and blood pressure data following the Klemera and Doubal method (KDM) (Klemera and Doubal, 2006) and the procedure described by Levine (Levine, 2013). KDM Biological Age details are reported in the **Supplemental Methods**. Participants with faster DunedinPoAm had more advanced KDM Biological Age (r=0.20 95% CI [0.15-0.26], p<0.001;) and worse self-rated health (r=−0.22 [−0.28,−0.16], p<0.001;). Covariate adjustment to models for estimated cell counts (Houseman et al., 2012) and smoking status did not change results. Results for all models are reported in **Supplemental Table 4**.

In comparison to DunedinPoAm, effect-sizes for associations with self-rated health and KDM Biological Age were smaller for the epigenetic clocks and, in the cases of the Horvath and Hannum clocks, were not statistically different from zero at the alpha=0.05 threshold (**Figure 3 Panels C and D**). Effect-sizes are reported in **Supplemental Table 4** and plotted **Supplemental Figure 3**.

### DunedinPoAm was associated with chronic disease morbidity and increased risk of mortality among older men in the Normative Aging Study (NAS)

To test if faster DunedinPoAm was associated with morbidity and mortality, we analyzed data from N=771 older men in the Veterans Health Administration Normative Aging Study (NAS; at baseline, mean chronological age=77, SD=7).

We first tested if higher DunedinPoAm levels, which indicate faster aging, were associated with increased risk of mortality. During follow-up from 1999-2013, 46% of NAS participants died over a mean follow-up of 7 years (SD=7). Those with faster DunedinPoAm at baseline were at increased risk of death (Hazard Ratio (HR)=1.29 [1.16-1.45], p<0.001; **Figure 4**).

**Figure 4.**
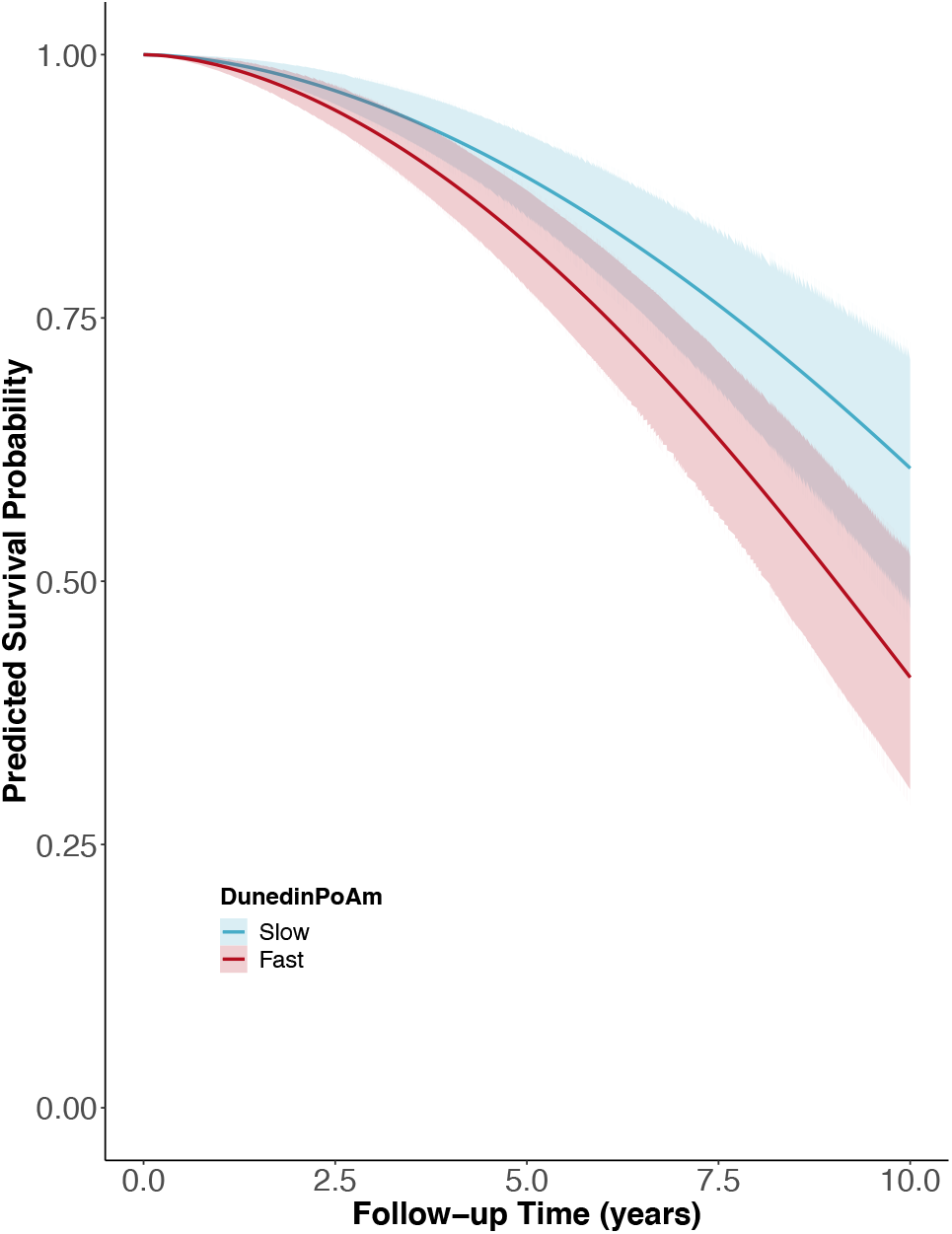
Association of DunedinPoAm with mortality in the Normative Aging Study. The figure plots predicted survival curves for participants with fast DunedinPoAm (1 SD or more above the mean; red shading) and slow DunedinPoAm (1 SD or more below the mean; blue shading). Survival curves were estimated based on Cox proportional hazard regressions of survival time on DunedinPoAm. Models included covariate adjustment for chronological age.

We next tested if NAS participants with faster DunedinPoAm experienced higher levels of chronic disease morbidity, measured as the count of diagnosed diseases (hypertension, type-2 diabetes, cardiovascular disease, chronic obstructive pulmonary disease, chronic kidney disease, and cancer). During follow-up across 4 assessments during 1999-2013 (n=1,448 observations of the N=771 participants), n=175 NAS participants were diagnosed with a new chronic disease. Those with faster baseline DunedinPoAm were at increased risk of new diagnosis (HR=1.19 [1.03-1.38], p<0.019). In repeated-measures analysis of prevalent chronic disease, faster DunedinPoAm was associated with having a higher level of chronic disease morbidity (IRR=1.15 [1.11-1.20], p<0.001).

Finally, we utilized the repeated-measures data to test if NAS participants’ DunedinPoAms increased as they aged. We tested within-person change in DuneidnPoAm over time (n=1,253 observations of N=536 participants with 2-4 timepoints of DNA methylation data). Consistent with Understanding Society analysis showing faster DunedinPoAm in older as compared to younger adults, NAS participants’ DunedinPoAms increased across repeated assessments. For every five years of follow-up, participants’ DunedinPoAms increased by 0.012 (SE=0.002) units, or about 0.2 standard deviations.

Covariate adjustment to models for estimated cell counts (Houseman et al., 2012) and smoking status did not change results, with the exception that the effect-size for DunedinPoAm was attenuated below the alpha=0.05 threshold of statistical significance in smoking-adjusted analysis of chronic disease incidence. Results for all models are reported in **Supplemental Table 5**.

In comparison to DunedinPoAm, effect-sizes for associations with mortality and chronic disease were smaller for the epigenetic clocks and were not statistically different from zero in many of the models (**Supplemental Table 5** and **Supplemental Figure 4**).

### Childhood exposure to poverty and victimization were associated with faster DunedinPoAm in young adults in the E-Risk Study

To test if DunedinPoAm indicated faster aging in young people with histories of exposure thought to shorten healthy lifespan, we analyzed data from N=1,658 members of the E-Risk Longitudinal Study. The E-Risk Study follows a 1994-95 birth cohort of same-sex twins. Blood DNA methylation data were collected when participants were aged 18 years. We analyzed two exposures associated with shorter healthy lifespan, childhood low socioeconomic status and childhood victimization. Socioeconomic status was measured from data on their parents’ education, occupation, and income (Trzesniewski et al., 2006). Victimization was measured from exposure dossiers compiled from interviews with the children’s mothers and home-visit assessments conducted when the children were aged 5, 7, 10, and 12 (Fisher et al., 2015). The dossiers recorded children’s exposure to domestic violence, peer bullying, physical and sexual harm by an adult, and neglect. 72% of the analysis sample had no victimization exposure, 21% had one type of victimization exposure, 4% had two types of exposure, and 2% had three or more types of exposure.

E-Risk adolescents who grew up in lower socioeconomic-status families exhibited faster DunedinPoAm (Cohen’s d for comparison of low to moderate SES =0.21 [0.06-0.35]; Cohen’s d for comparison of low to high SES =0.44 [0.31-0.56]; Pearson r=0.19 [0.13-0.24]). In parallel, E-Risk adolescents with exposure to more types of victimization exhibited faster DunedinPoAm (Cohen’s d for comparison of never victimized to one type of victimization =0.28 [0.15-0.41]; Cohen’s d for comparison of never victimized to two types of victimization =0.48 [0.23-0.72]; Cohen’s d for comparison of never victimized to three or more types of victimization =0.53 [0.25-0.81]; Pearson r=0.15 [0.10-0.20]). Covariate adjustment to models for estimated cell counts (Houseman et al., 2012) did not change results. Adjustment for smoking status attenuated effect-sizes by about half, but most associations remained statistically different from zero at the alpha=0.05 level. Results for all models are reported in **Supplemental Table 6**. Differences in DunedinPoAm across strata of childhood socioeconomic status and victimization are graphed in **Figure 5**.

**Figure 5.**
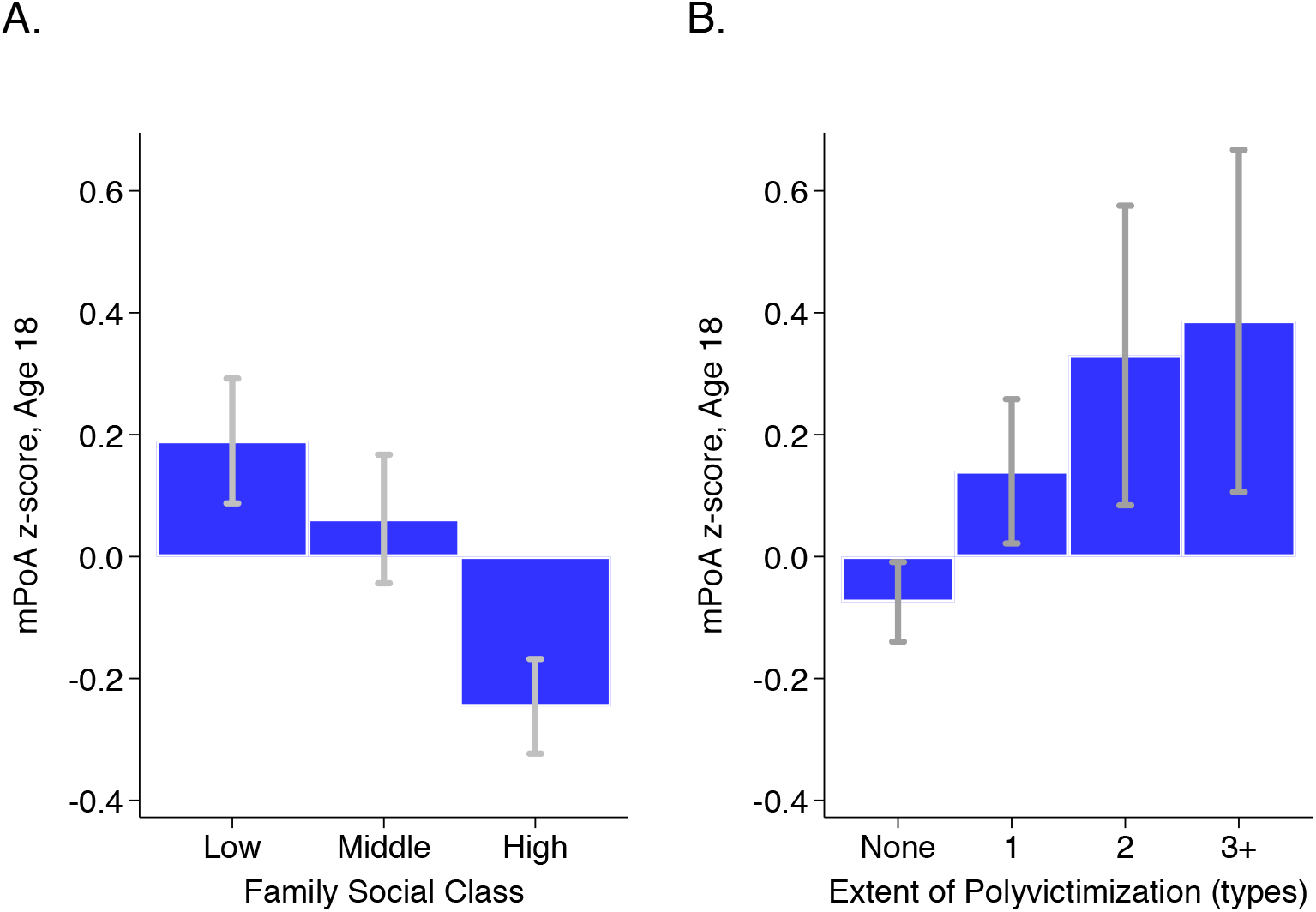
DunedinPoAm levels by strata of childhood socioeconomic status (SES) and victimization in the E-Risk Study. Panel A (left side) plots means and 95% CIs for DunedinPoAm measured at age 18 among E-Risk participants who grew up low, middle, and high socioeconomic status households. Panel B (right side) plots means and 95% CIs for DunedinPoAm measured at age 18 among E-Risk participants who experienced 0, 1, 2, or 3 or more types of victimization through age 12 years.

In comparison to DunedinPoAm, effect-sizes for associations with childhood socioeconomic circumstances and victimization were smaller for the epigenetic clocks and, in the cases of the Horvath and Hannum clocks, were not statistically different from zero at the alpha=0.05 threshold. Effect-sizes are reported in **Supplemental Table 6** and plotted in **Supplemental Figure 5**.

### DunedinPoAm measured at baseline in the CALERIE randomized trial predicted future rate of aging measured from clinical-biomarker data

The CALERIE Trial is the first randomized trial of long-term caloric restriction in non-obese adult humans. CALERIE randomized N=220 adults on a 2:1 ratio to treatment of 25% caloric restriction (CR-treatment) or control ad-libitum (AL-control, as usual) diet for two years (Ravussin et al., 2015). We previously reported that CALERIE participants who were randomized to CR-treatment experienced a slower rate of biological aging as compared to participants in the AL-control arm based on longitudinal change analysis of clinical-biomarker data from the baseline, 12-month, and 24-month follow-up assessments (Belsky et al., 2017b). Among control participants, the rate of increase in biological age measured using the Klemera-Doubal method (KDM) Biological Age algorithm was 0.71 years of biological age per 12-month follow-up interval. (This slower-than-expected rate of aging could reflect differences between CALERIE Trial participants, who were selected for being in good health, and the nationally representative NHANES sample in which the KDM algorithm was developed (Belsky et al., 2017b).) In contrast, among treatment participants, the rate of increase was only 0.11 years of biological age per 12-month follow-up interval (difference b=−0.60 [−0.99, −0.21]). We subsequently generated DNA methylation data from blood DNA that was collected at the baseline assessment of the CALERIE trial for a sub-sample (N=68 AL-control participants and 118 CR-treatment participants). We used these methylation data to calculate participants’ DunedinPoAm values at study baseline. We then tested if baseline DunedinPoAm could predict participants’ future rate of biological aging as they progressed through the trial.

We first replicated our original analysis within the methylation sub-sample. Results were the same as in the full sample (**Supplemental Table 7**). Next, we compared DunedinPoAm between CR-treatment and AL-control participants. As expected, there was no group difference at baseline (AL M=1.00, SD=0.05; CR M=1.01, SD=0.06, p-value for difference =0.440). Finally, we tested if participants’ baseline DunedinPoAm was associated with their rate of biological aging over the 24 months of follow-up, and if this association was modified by randomization to caloric restriction as compared to ad libitum diet. For AL-control participants, faster baseline DunedinPoAm predicted faster biological aging over the 24-month follow-up, although in this small group this association was not statistically significant at the alpha=0.05 level (b=0.22 [−0.05, 0.49], p=0.104). For CR-treatment participants, the association of baseline DunedinPoAm with future rate of aging was sharply reduced, (b=−0.08 [−0.24, 0.09], p=0.351), although the difference between the rate of aging in the AL-control and CR-treatment groups did not reach the alpha=0.05 threshold for statistical significance (interaction-term testing difference in slopes b=−0.30 [−0.61, 0.01], p-value=0.060). Slopes of change in KDM Biological Age for participants in the AL-control and CR-treatment groups are plotted for fast baseline DunedinPoAm (1 SD above the mean) and slow baseline DunedinPoAm (1 SD below the mean) in **Figure 6**. CALERIE DNA methylation data are not yet available to test if the intervention altered post-treatment DunedinPoAm.

**Figure 6.**
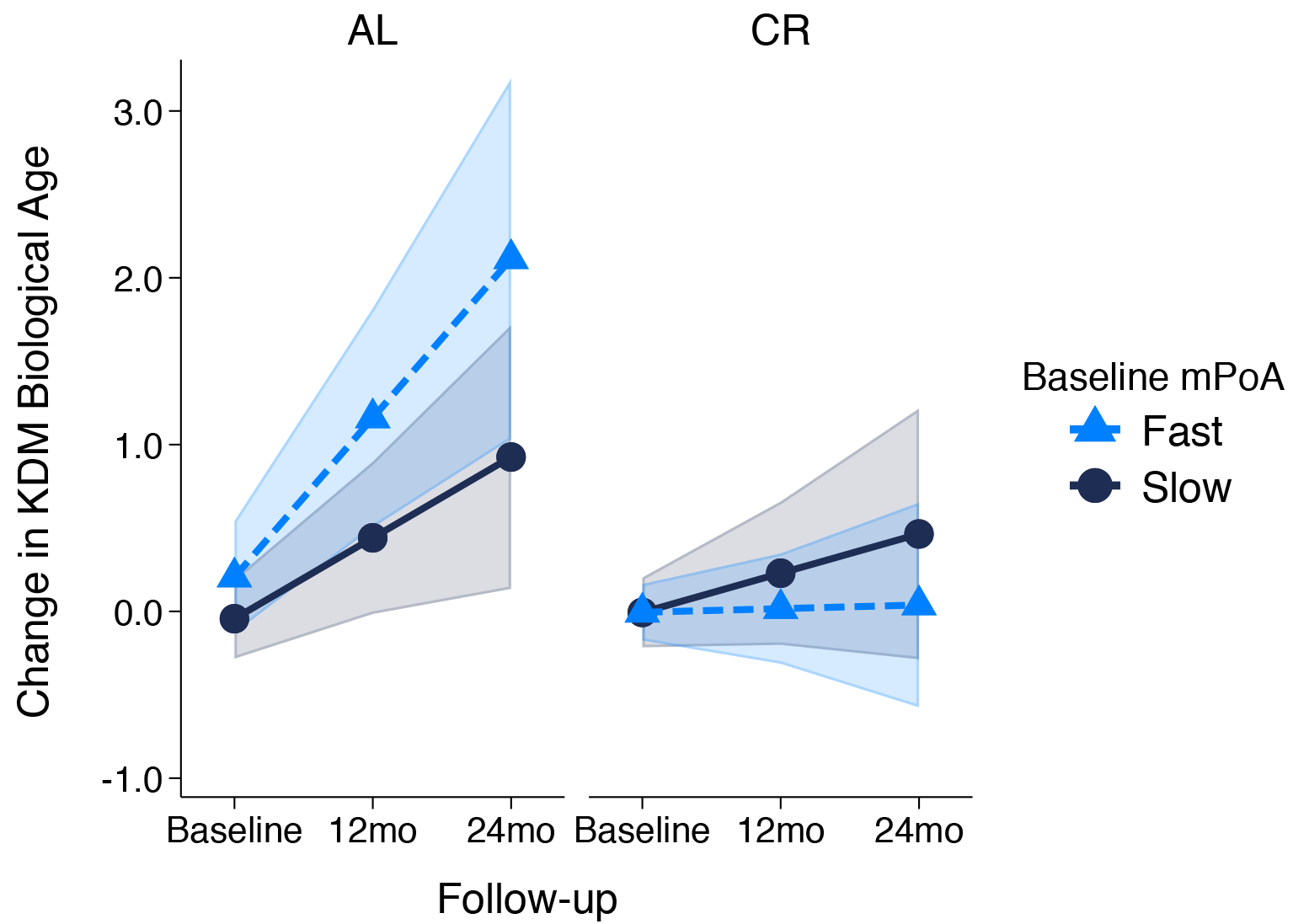
Change in KDM Biological Age over 24-month follow-up by treatment condition and baseline DunedinPoAm in the CALERIE Trial. Fast DunedinPoAm is defined as 1 SD above the sample mean. Slow DunedinPoAm is defined as 1 SD below the cohort mean. Slopes are predicted values from mixed effects regression including a 3-way interaction between trial condition, time, and continuous DunedinPoAm at baseline. The figure shows that in the Ad Libitum (AL) arm of the trial, participants with fast DunedinPoAm at baseline experience substantially more change in KDM Biological Age from baseline to follow-up as compared to AL participants with slow DunedinPoAm. In contrast, there was little difference between participants with fast as compared to slow DunedinPoAm in the Caloric Restriction (CR) arm of the trial.

## DISCUSSION

Breakthrough discoveries in the new field of geroscience suggest opportunities to extend healthy lifespan through interventions that slow biological processes of aging (Campisi et al., 2019). To advance translation of these interventions, measures are needed that can detect changes in a person’s rate of biological aging (Moffitt et al., 2016). We previously showed that the rate of biological aging can be measured by tracking change over time in multiple indicators of organ-system integrity (Belsky et al., 2015). Here, we report data illustrating the potential to streamline measurement of Pace of Aging to an exportable, inexpensive and non-invasive blood test, and thereby ease implementation of Pace of Aging measurement in studies of interventions to slow processes of biological aging.

We conducted machine-learning analysis of the original Pace of Aging measure using elastic-net regression and whole-genome blood DNA methylation data. We trained the algorithm to predict how fast a person was aging. We called the resulting algorithm “DunedinPoAm” for “(m)ethylation (P)ace (o)f (A)ging”. There were four overall findings:

First, while DunedinPoAm was not a perfect proxy of Pace of Aging, it nevertheless captured critical information about Dunedin Study members’ healthspan-related characteristics. Across the domains of physical function, cognitive function, and subjective signs of aging, Study members with faster DunedinPoAm at age 38 were worse off seven years later at age 45 and, in repeated-measures analysis of change, they showed signs of more rapid decline. Effect-sizes were equal to or greater than those for the 18-biomarker 3-time point measure of Pace of Aging. This result suggests that the DNA-methylation elastic-net regression used to develop DunedinPoAm may have distilled the aging signal from the original Pace of Aging measure and excluded some noise. In sum, DunedinPoAm showed promise as an easy-to-implement alternative to Pace of Aging. Emerging technologies for deep-learning analysis (Zhavoronkov et al., 2019) may improve methylation measurement of Pace of Aging. Alternatively, integration of methylation data with additional molecular datasets (Hasin et al., 2017; Zierer et al., 2015) may be needed to achieve precise measurement of Pace of Aging from a single time-point blood sample.

Second, DunedinPoAm analysis of the Understanding Society and Normative Aging Study samples provided proof-of-concept for using DunedinPoAm to quantify biological aging. Age differences in DunedinPoAm parallel population demographic patterns of mortality risk. In the Understanding Society sample, older adults had faster DunedinPoAm as compared to younger ones. In the Normative Aging Study sample, participants’ DunedinPoAm values increased as they aged. These observations are consistent with the well-documented acceleration of mortality risk with advancing chronological age (Robine, 2011). However, it sets DunedinPoAm apart from other indices of biological aging, which are not known to register this acceleration (Finch and Crimmins, 2016; Li et al., 2020). DunedinPoAm may therefore provide a novel tool for testing how the rate of aging changes across the life course and whether, as demographic data documenting so-called “mortality plateaus” suggest, processes of aging slow down at the oldest chronological ages (Barbi et al., 2018).

DunedinPoAm is related to but distinct from alternative approaches to quantification of biological aging. DunedinPoAm was moderately correlated with aging rates measured by the epigenetic clocks proposed by Hannum et al. and Levine et al. (Hannum et al., 2013; Levine et al., 2018) as well as KDM Biological Age derived from clinical biomarker data (Klemera and Doubal, 2006; Levine, 2013), and with self-rated health. Consistent with findings for the measured Pace of Aging (Belsky et al., 2018), DunedinPoAm was only weakly correlated with the multi-tissue clock proposed by Horvath. DunedinPoAmwas more strongly correlated with a clinical-biomarker measure of biological age, with self-rated health, with functional test-performance and decline, and with morbidity and mortality as compared to the epigenetic clocks.

Third, DunedinPoAm is already variable by young adulthood and is accelerated in young people at risk for eventual shortened healthspan. E-Risk young adults who grew up in socioeconomically disadvantaged families or who were exposed to victimization early in life already showed accelerated DunedinPoAm by age 18, consistent with epidemiological observations of shorter healthy lifespan for individuals with these exposures (Adler and Rehkopf, 2008; Danese and McEwen, 2012). We previously found that Dunedin Study members with histories of early-life adversity showed accelerated Pace of Aging in their 30s (Belsky et al., 2017a). DunedinPoAm analysis of the E-Risk cohort suggests effects may be already manifest at least a decade earlier. DunedinPoAm may therefore provide a useful index that can be applied to evaluate prevention programs to buffer at-risk youth against health damaging effects of challenging circumstances.

Fourth, DunedinPoAm analysis of the CALERIE trial provided proof-of-concept for using DunedinPoAm to quantify biological aging in geroprotector intervention studies. DunedinPoAm measures the rate of aging over the recent past. Control-arm participants’ baseline DunedinPoAm correlated positively with their clinical-biomarker pace of aging over the two years of the trial, consistent with the hypothesis that their rate of aging was not altered. In contrast, there was no relationship between DunedinPoAm and clinical-biomarker pace of aging for caloric-restriction-arm participants, consistent with the hypothesis that caloric restriction altered participants’ rate of aging. Ultimately, data on DunedinPoAm for all CALERIE participants (and participants in other geroprotector trails) at trial baseline and follow-up will be needed to establish utility of DunedinPoAm as a surrogate endpoint. In the mean-time, these data establish potential to use DunedinPoAm as a pre-treatment covariate in geroprotector trials to boost statistical power (Kahan et al., 2014) or to screen participants for enrollment, e.g. to identify those who are aging more rapidly and may therefore show larger effects of treatment.

We acknowledge limitations. Foremost, DunedinPoAm is a first step toward a single-assay cross-sectional measurement of Pace of Aging. The relatively modest size of the Dunedin cohort and the lack of other cohorts that have the requisite 3 or more waves of repeated biomarkers to measure the Pace of Aging limited sample size for our machine-learning analysis to develop methylation algorithms. As Pace of Aging is measured in additional cohorts, more refined analysis to develop DunedinPoAm-type algorithms will become possible. In addition, our work thus far has not addressed population diversity in biological aging. The Dunedin cohort in which DunedinPoAm was developed and the Understanding Society and E-Risk cohorts and CALERIE trial sample in which it was tested were mostly of white European-descent. Follow-up of DunedinPoAm in more diverse samples is needed to establish cross-population validity. Finally, because methylation data are not yet available from CALERIE follow-up assessments, we could not test if intervention modified DunedinPoAm at outcome. Ultimately, to establish DunedinPoAm as a surrogate endpoint for healthspan, it will be necessary to establish not only robust association with healthy lifespan phenotypes and modifiability by intervention, but also the extent to which changes in DunedinPoAm induced by intervention correspond to changes in healthy-lifespan phenotypes (Prentice, 1989).

Within the bounds of these limitations, our analysis establishes proof-of-concept for DunedinPoAm as a single-time-point measure that quantifies Pace of Aging from a blood test. It can be implemented in Illumina 450k and EPIC array data, making it immediately available for testing in a wide range of existing datasets as a complement to existing methylation measures of aging. Critically, DunedinPoAm offers a unique measurement for intervention trials and natural experiment studies investigating how the rate of aging may be changed by behavioral or drug therapy, or by environmental modification. DunedinPoAm may be especially valuable to studies that collect data outside of clinical settings and lack blood chemistry, hematology, and other data needed to measure aging-related changes to physiology.

## Acknowledgement

This research was supported by US-National Institute on Aging grants AG032282and UK Medical Research Council grant MR/P005918/1. The Dunedin Multidisciplinary Health and Development Research Unit is supported by the New Zealand Health Research Council Programme Grant (16-604), and the New Zealand Ministry of Business, Innovation and Employment (MBIE). We thank the Dunedin Study members, Unit research staff, and Study founder Phil Silva.

Understanding Society data come from The UK Household Longitudinal Study, which is led by the Institute for Social and Economic Research at the University of Essex and funded by the Economic and Social Research Council (ES/M008592/1). The data were collected by NatCen and the genome wide scan data were analysed by the Wellcome Trust Sanger Institute. Information on how to access the data can be found on the Understanding Society website https://www.understandingsociety.ac.uk/. Data governance was provided by the METADAC data access committee, funded by ESRC, Wellcome, and MRC (2015-2018: MR/N01104X/1; 2018-2020: ES/S008349/1)

The Normative Aging Study is supported by the National Institute of Environmental Health Sciences (grants P30ES009089, R01ES021733, R01ES025225, and R01ES027747). The VA Normative Aging Study is supported by the Cooperative Studies Program/Epidemiology Research and Information Center of the U.S. Department of Veterans Affairs and is a component of the Massachusetts Veterans Epidemiology Research and Information Center, Boston, Massachusetts.

The E-Risk Study is supported by the UK Medical Research Council (grant G1002190), the US National Institute of Child Health and Development (grant HD077482), and the Jacobs Foundation. The generation of DNA methylation data was supported by the American Asthma Foundation.

This investigation was made possible in part through use of the CALERIE data repository and was supported in part by U24AG047121 and R01AG061378.

This work used a high-performance computing facility partially supported by grant 2016-IDG-1013 (HARDAC+: Reproducible HPC for Next-generation Genomics”) from the North Carolina Biotechnology Center.

DWB was additionally supported by US National Institute on Aging grant R21AG054846 and the Jacobs Foundation.

## SUPPLEMENTAL MATERIALS

### 1. Data

**The Dunedin Study** is a longitudinal investigation of health and behavior in a complete birth cohort. Study members (N=1,037; 91% of eligible births; 52% male) were all individuals born between April 1972 and March 1973 in Dunedin, New Zealand (NZ), who were eligible based on residence in the province and who participated in the first assessment at age 3. The cohort represents the full range of socioeconomic status on NZ’s South Island and matches the NZ National Health and Nutrition Survey on key health indicators (e.g., BMI, smoking, GP visits) (Poulton et al., 2015). The cohort is primarily white (93%) (Poulton et al., 2015). Assessments were carried out at birth and ages 3, 5, 7, 9, 11, 13, 15, 18, 21, 26, 32, 38 and, most recently, 45 years, when 94% of the 997 study members still alive took part. At each assessment, each study member is brought to the research unit for a full day of interviews and examinations. Study data may be accessed through agreement with the Study investigators (https://moffittcaspi.trinity.duke.edu/research-topics/dunedin).

Dunedin Study analysis to develop the Pace of Aging algorithm included 18 biomarkers measured at the age 26, 32, and 38 assessments: (in order of listing in Figure 3 of (Belsky et al., 2015)) glycated hemoglobin, cardiorespiratory fitness, waist-hip ratio, FEV_1_/FVC ratio, FEV_1_, mean arterial pressure, body mass index, leukocyte telomere length, creatinine clearance, blood urea nitrogen, lipoprotein (a), triglycerides, gum health, total cholesterol, white blood cell count, high-sensitivity C-reactive protein, HDL cholesterol, ApoB100/ApoA1 ratio.

**Supplemental Table 1.** Physical and cognitive functioning and subjective signs of aging measures in the Dunedin Study.

| Physical Functioning (N=800 with DunedinPoAm data) |  |
| --- | --- |
| Balance | Balance was measured using the Unipedal Stance Test as the maximum time achieved across three trials of the test with eyes closed (Bohannon et al., 1984; Springer et al., 2007; Vereeck et al., 2008). |
| Gait Speed | Gait speed (meters per second) was assessed with the 6-m-long GAITRite Electronic Walkway (CIR Systems, Inc) with 2-m acceleration and 2-m deceleration before and after the walkway, respectively. Gait speed was assessed under 3 walking conditions: usual gait speed (walk at normal pace from a standing start, measured as a mean of 2 walks) and 2 challenge paradigms, dual-task gait speed (walk at normal pace while reciting alternate letters of the alphabet out loud, starting with the letter "A," measured as a mean of 2 walks) and maximum gait speed (walk as fast as safely possible, measured as a mean of 3 walks). We calculated the mean of the 3 individual walk conditions to generate our primary measure of composite gait speed (Rasmussen et al., 2019). |
| Steps in Place | The 2-min step test was measured as the number of times a participant lifted their right knee to mid-thigh height (measured as the height half-way between the knee cap and the iliac crest) in 2 minutes at a self-directed pace (Jones and Rikli, 2002; Rikli and Jones, 1999). |
| Chair Stands | Chair rises were measured as the number of stands a participant completed in 30 seconds from a seated position (Jones et al., 1999; Jones and Rikli, 2002). |
| Grip Strength | Handgrip strength was measured for the dominant hand (elbow held at 90°, upper arm held tight against the trunk) as the maximum value achieved across three trials using a Jamar digital dynamometer (Mathiowetz et al., 1985; Rantanen T et al., 1999). |
| Motor Coordination | At ages 38 and 45, we measured motor functioning as the time to completion of the Grooved Pegboard Test with the dominant hand. |
| Physical Limitations | Physical limitations were measured with the 10-item RAND 36-Item Health Survey 1.0 physical functioning scale (Ware and Sherbourne, 1992). Participant responses (“limited a lot”, “limited a little”, “not limited at all”) assessed their difficulty with completing various activities, e.g., climbing several flights of stairs, walking more than 1 km, participating in strenuous sports, etc. Scores were reversed to reflect physical limitations so that a high score indicates more limitations. |
| Decline in Physical Functioning | Tests of balance and grip strength and interviews about physical limitations were completed at both the age-38 and age-45 Dunedin Study assessments. We measured decline across the 7-year measurement interval by subtracting the age-38 test score from the age-45 test score. |
| **Cognitive Functioning (N=795 with mPoA data)** |  |
| Cognitive Functioning | The Wechsler Adult Intelligence Scale-IV (WAIS-IV) (Wechsler, 2008) was administered to the participants at age 45 years, yielding the IQ. In addition to full scale IQ, the WAIS-IV measures four specific domains of cognitive function: Processing Speed, Working Memory, Perceptual Reasoning, and Verbal Comprehension. |
| Cognitive Decline | IQ is a highly reliable measure of general intellectual functioning that captures overall ability across differentiable cognitive functions. We measured IQ from the individually administered Wechsler Intelligence Scale for Children-Revised (WISC-R; averaged across ages 7, 9, 11, and 13)(Wechsler, 2003) and the Wechsler Adult Intelligence Scale-IV (WAIS-IV; age 45) (Wechsler, 2008). We measured IQ decline by comparing scores from the WISC-R and the WAIS-IV. |
| **Subjective Signs of Aging (N=802 with mPoA data)** |  |
| Self-rated Health | Study members rated their health on a scale of 1-5 (poor, fair, good, very good, or excellent). |
| Facial Aging | Facial Aging is the subjective perception of aged appearance based on a facial photograph and is proposed as a clinically-useful marker of mortality risk (Christensen et al., 2009). Facial Aging measurement in the Dunedin Study was based on ratings by an independent panel of 8 raters of each participant’s facial photograph (Belsky et al., 2015; Shalev et al., 2014). Facial Aging was based on two measurements of perceived age. First, Age Range was assessed by an independent panel of 4 raters, who were presented with standardized (non-smiling) facial photographs of participants and were kept blind to their actual age. Raters used a Likert scale to categorize each participant into a 5- |
|  | year age range (i.e., from 20-24 years old up to 70+ years old) (interrater reliability = .77). Scores for each participant were averaged across all raters. Second, Relative Age was assessed by a different panel of 4 raters, who were told that all photos were of people aged 45 years old. Raters then used a 7-item Likert scale to assign a “relative age” to each participant (1=“young looking”, 7=“old looking”) (interrater reliability = .79). The measure of perceived age at 45 years, Facial Age, was derived by standardizing and averaging Age Range and Relative Age scores. |
| Subjective Decline | Self-rated Health and Facial Aging were measured at both the age-38 and age-45 assessments. We measured decline in self-rated health as incident fair/poor health reported at the age-45 assessment. We measured acceleration in Facial Aging by computing the difference in Facial Aging Z-scores between the age-45 and age-38 assessments. |

**Understanding Society** is an ongoing panel study of the United Kingdom population (https://www.understandingsociety.ac.uk/). During 2010-12, participants were invited to take part in a nurse’s exam involving a blood draw. Of the roughly 20,000 participants who provided clinical data in this exam, methylation data have been generated for just under 1,200. We analyzed data from 1,175 participants with available methylation and blood chemistry data. Documentation of the methylation (University of Essex, n.d.) and blood chemistry (University of Essex, n.d.) data resource is available online (https://www.understandingsociety.ac.uk/sites/default/files/downloads/documentation/health/user-guides/7251-UnderstandingSociety-Biomarker-UserGuide-2014.pdf).

#### Klemera-Doubal method (KDM) Biological Age

We measured KDM Biological age from blood chemistry, systolic blood pressure, and lung-function data using the algorithm proposed by Klemera and Doubal (Klemera and Doubal, 2006) trained in data from the NHANES following the method originally described by Levine (Levine, 2013) and using the dataset compiled by Hastings (Hastings et al., 2019). We included 8 of Levine’s original 10 biomarkers in the algorithm: albumin, alkaline phosphatase (log), blood urea nitrogen, creatinine (log), C-reactive protein (log), HbA1C, systolic blood pressure, and forced expiratory volume in 1 second (FEV_1_). We omitted total cholesterol because of evidence this biomarker shows different directions of association with aging in younger and older adults (Arbeev et al., 2016). Cytomegalovirus optical density was not available in the Understanding Society database.

#### Self-rated Health

Understanding Society participants rated their health as excellent, very-good, good, fair, or poor. We standardized this measure to have Mean=0, Standard Deviation=1 for analysis.

**The Normative Aging Study (NAS)** is an ongoing longitudinal study on aging established by the US Department of Veterans Affairs in 1963. Details of the study have been published previously (Bell et al., 1972). Briefly, the NAS is a closed cohort of 2,280 male veterans from the Greater Boston area enrolled after an initial health screening to determine that they were free of known chronic medical conditions. Participants have been re-evaluated every 3–5 years on a continuous rolling basis using detailed on-site physical examinations and questionnaires. DNA from blood samples was collected from 771 participants beginning in 1999. We analyzed blood DNA methylation data from up to four repeated assessments conducted through 2013 (Gao et al., 2019a; Panni Tommaso et al., 2016). Of the 771 participants with DNA methylation data, n=536 (46%) had data from 2 repeated assessments and n=178 (23%) had data from three or four repeated assessments. We restricted the current analysis to participants with at least one DNA methylation data point. The NAS was approved by the Department of Veterans Affairs Boston Healthcare System and written informed consent was obtained from each subject before participation.

#### Mortality

Regular mailings to study participants have been used to acquire vital-status information and official death certificates were obtained from the appropriate state health department to be reviewed by a physician. Participant deaths are routinely updated by the research team and the last available update was on 31 December 2013.. During follow-up, n=355 (46%) of the 771 NAS participants died.

#### Chronic Disease Morbidity

We measured chronic disease morbidity from participants medical histories and prior diagnoses (Gao et al., 2019b, 2019c; Lepeule et al., 2018; Nyhan et al., 2018). We counted the number of chronic diseases to compose an ordinal index with categories of 0, 1, 2, 3, or 4+ of the following comorbidities: hypertension, type-2 diabetes, cardiovascular disease, chronic obstructive pulmonary disease, chronic kidney disease, and cancer.

**The Environmental Risk Longitudinal Twin Study** tracks the development of a birth cohort of 2,232 British participants. The sample was drawn from a larger birth register of twins born in England and Wales in 1994-1995. Full details about the sample are reported elsewhere (Moffitt and E-risk Team, 2002). Briefly, the E-Risk sample was constructed in 1999-2000, when 1,116 families (93% of those eligible) with same-sex 5-year-old twins participated in home-visit assessments. This sample comprised 56% monozygotic (MZ) and 44% dizygotic (DZ) twin pairs; sex was evenly distributed within zygosity (49% male). Families were recruited to represent the UK population of families with newborns in the 1990s, on the basis of residential location throughout England and Wales and mother’s age. Teenaged mothers with twins were over-selected to replace high-risk families who were selectively lost to the register through non-response. Older mothers having twins via assisted reproduction were under-selected to avoid an excess of well-educated older mothers. The study sample represents the full range of socioeconomic conditions in the UK, as reflected in the families’ distribution on a neighborhood-level socioeconomic index (ACORN [A Classification of Residential Neighborhoods], developed by CACI Inc. for commercial use): 25.6% of E-Risk families lived in “wealthy achiever” neighborhoods compared to 25.3% nationwide; 5.3% vs. 11.6% lived in “urban prosperity” neighborhoods; 29.6% vs. 26.9% lived in “comfortably off” neighborhoods; 13.4% vs. 13.9% lived in “moderate means” neighborhoods, and 26.1% vs. 20.7% lived in “hard-pressed” neighborhoods. E-Risk underrepresents “urban prosperity” neighborhoods because such households are likely to be childless.

Home-visits assessments took place when participants were aged 5, 7, 10, 12 and, most recently, 18 years, when 93% of the participants took part. At ages 5, 7, 10, and 12 years,assessments were carried out with participants as well as their mothers (or primary caretakers); the home visit at age 18 included interviews only with participants. Each twin was assessed by a different interviewer. These data are supplemented by searches of official records and by questionnaires that are mailed, as developmentally appropriate, to teachers, and co-informants nominated by participants themselves. The Joint South London and Maudsley and the Institute of Psychiatry Research Ethics Committee approved each phase of the study. Parents gave informed consent and twins gave assent between 5-12 years and then informed consent at age 18. Study data may be accessed through agreement with the Study investigators (https://moffittcaspi.trinity.duke.edu/research-topics/erisk).

#### Childhood Socioeconomic Status

(SES) was defined through a standardized composite of parental income, education, and occupation (Trzesniewski et al., 2006). The three SES indicators were highly correlated (r=0.57–0.67) and loaded significantly onto one factor. The population-wide distribution of the resulting factor was divided in tertiles for analyses.

#### Childhood Victimization

As previously described (Danese et al., 2016), we assessed exposure to six types of childhood victimization between birth to age 12: exposure to domestic violence between the mother and her partner, frequent bullying by peers, physical and sexual harm by an adult, and neglect.

##### CALERIE

The CALERIE trial is described in detail elsewhere (Ravussin et al., 2015). Briefly, N=220 normal-weight (22.0 ≤ BMI < 28 kg/m^2^) participants (70% female, 77% white) aged 21-50 years at baseline were randomized to caloric restriction or ad libitum conditions with a 2:1 ratio (n=145 to caloric restriction, n=75 to ad libitum). “*Ad libitum*” (normal) caloric intake was determined from two consecutive 14-day assessments of total daily energy expenditure using doubly labeled water (Redman et al., 2014). Average percent caloric restriction over six-month intervals was retrospectively calculated by the intake-balance method with simultaneous measurements of total daily energy expenditure using doubly labeled water and changes in body composition (Racette et al., 2012; Wong et al., 2014). Over the course of the trial, participants in the caloric-restriction arm averaged 12% reduction in caloric intake (about half the prescribed reduction). Participants in the ad libitum condition reduced caloric intake by <2% (Ravussin et al., 2015). CALERIE data are available at https://calerie.duke.edu/samples-data-access-and-analysis.

#### Klemera-Doubal method (KDM) Biological Age

KDM Biological age was measured according to the procedure described in our previous article (Belsky et al., 2017). Briefly, we computed KDM Biological Age from CALERIE blood chemistry and blood pressure data using the algorithm proposed by Klemera and Doubal (Klemera and Doubal, 2006) trained in data from the NHANES following the method originally described by Levine (Levine, 2013) and NHANES data from years matched to the timing of the CALERIE Trial. We included 8 of Levine’s original 10 biomarkers in the algorithm: albumin, alkaline phosphatase (log), blood urea nitrogen, creatinine (log), C-reactive protein (log), HbA1C, systolic blood pressure, and total cholesterol. Cytomegalovirus optical density and lung function were not measured in CALERIE. We supplemented the algorithm with data on uric acid and white blood cell count.

### 2. Sensitivity analyses

We tested sensitivity of mPoA to alternative methods of normalizing DNA methylation data. We normalized data using the ‘methylumi’ and ‘minfi’ packages and computed correlations between mPoA measures derived from these two datasets. The correlation was r=0.94.

The elastic net model selected 46 CpGs to compose the mPoA. One of these CpGs, cg11897887, has been identified as an mQTL (Volkov et al., 2016). To evaluate sensitivity of results to the exclusion of this SNP, we computed a version of the mPoA excluding this CpG and repeated analysis. This version of the score was correlated with the full mPoA at r=1. Results were the same in analyses with both versions (available from the authors upon request).

Another CpG selected in the elastic net, cg05575921, is located within the gene *AHRR*, previously identified as a methylation site modified by tobacco exposure and associated with lung cancer and other chronic disease, e.g. (Fasanelli et al., 2015; Reynolds et al., 2015). We tested sensitivity of results to the exclusion of this probe using the method described above. This version of the score was correlated with the full mPoA at r=0.94. Again, results were the same in analyses with both versions (available from the authors upon request).

### 3. Bootstrap repetition analysis to estimate out-of-sample correlation between mPoA and longitudinal Pace of Aging

The Dunedin Study is the only dataset to include measured 12-year longitudinal Pace of Aging. To estimate the out-of-sample correlation between mPoA and the original Pace of Aging measure, we conducted 90/10 crossfold validation analysis. We randomly selected 90% of the cohort to serve as the “training” sample in which the mPoA algorithm was developed. We used the remaining 10% to form a “test” sample to estimate the correlation between mPoA and Pace of Aging. We repeated this analysis across 100 bootstrap repetitions. In each repetition, we randomly sampled 90% of the cohort to use in the training analysis and reserved the remaining 10% for testing.

The mPoA algorithms developed across the 100 bootstrap repetitions included different sets of CpGs (range of 21-209 CpGs selected, M=54, SD=27 CpGs). However, the resulting algorithms were highly correlated (mean pairwise r=0.90, SD=0.14). The average correlation between the 90%-trained mPoA and longitudinal Pace of Aging in the 10% test samples was (r=0.33, SD=0.10). Details are reported in **Supplemental Figure 2**.

---

**Supplemental Table 1. Probes and associated weights composing the DunedinPoAm algorithm.** The DunedinPoAm algorithm is a linear combination of 46 CpG methylation beta values weighted by coefficients estimated in the elastic net regression and added to the model intercept value of −0.06. (For sensitivity analyses addressing normalization method, and specific probes, see Supplement section 3.)

**Supplemental Table 2.** Comparison of age-38 DundedinPoAm and age 26-38 Pace of Aging effect-sizes for analysis of healthspan-related characteristics. The table shows effect-sizes for analysis of healthspan-related characteristics at age 45 years. mPoA was measured from blood DNA methylation collected when Study members were aged 38 years. Pace of Aging was measured from longitudinal change in 18 biomarkers across measurements made at ages 26, 32, and 38 years. Sample restricted to N=810 Study members with data on mPoA and Pace of Aging. Effect-sizes correspond to the analysis reported in **Supplemental Figure 1B** and are reported in terms of standard deviation differences in the age-45 outcome associated with a 1 standard deviation increase in mPoA (i.e. effect-sizes are interpretable as Pearson r). All models were adjusted for sex.

|  | DunedinPoAm Effect-Sizes |  |  | Pace of Aging Effect-Sizes |  |  |
| --- | --- | --- | --- | --- | --- | --- |
|  | r | 95% CI | p-value | r | 95% CI | p-value |
| Balance | -0.29 | [-0.36, -0.23] | 5.13E-18 | -0.21 | [-0.27, -0.15] | 9.32E-11 |
| Gait Speed | -0.27 | [-0.33, -0.20] | 2.15E-14 | -0.18 | [-0.25, -0.11] | 4.77E-07 |
| Steps in Place | -0.22 | [-0.29, -0.15] | 1.70E-09 | -0.16 | [-0.23, -0.08] | 3.98E-05 |
| Chair Stands | -0.19 | [-0.26, -0.12] | 8.96E-08 | -0.18 | [-0.25, -0.11] | 5.01E-07 |
| Grip Strength | -0.05 | [-0.12, 0.02] | 0.138 | -0.07 | [-0.14, 0.00] | 0.060 |
| Motor Coordination | -0.20 | [-0.27, -0.13] | 5.06E-08 | -0.18 | [-0.25, -0.11] | 7.10E-07 |
| Physical Limitations | 0.21 | [0.12, 0.30] | 3.08E-06 | 0.15 | [0.07, 0.22] | 1.77E-04 |
| Perceptual Reasoning | -0.29 | [-0.35, -0.22] | 2.51E-17 | -0.19 | [-0.26, -0.11] | 6.60E-07 |
| Working Memory | -0.21 | [-0.28, -0.15] | 4.35E-10 | -0.16 | [-0.23, -0.10] | 1.49E-06 |
| Processing Speed | -0.15 | [-0.22, -0.09] | 6.86E-06 | -0.16 | [-0.23, -0.08] | 3.46E-05 |
| Self-rated Health | -0.28 | [-0.35, -0.20] | 1.86E-13 | -0.24 | [-0.31, -0.16] | 6.55E-10 |
| Facial Aging | 0.35 | [0.28, 0.43] | 9.94E-19 | 0.26 | [0.19, 0.34] | 1.34E-11 |

**Supplemental Table 3.** Effect-sizes for associations of age-38 DundedinPoAm and epigenetic clocks with healthspan-related characteristics at age 45 and change in healthspan characteristics from age 38-45. Panel A of the table shows effect-sizes for analysis of healthspan-related characteristics at age 45 years. Panel B of the table shows effect-sizes for analysis of change in healthspan-related characteristics from 38 to 45. Methylation measurements were derived from blood DNA methylation collected when Study members were aged 38 years. Prior to analysis, Horvath, Hannum, and Levine Clock values were residualized for chronological age. Effect-sizes are reported in terms of standard deviation differences in the outcome associated with a 1 standard deviation increase in methylation measures (i.e. effect-sizes are interpretable as Pearson r). All models were adjusted for sex.

**Panel A. Analysis of healthspan-related characteristics at age 45 years**
|  | DunedinPoAm |  |  | Horvath Clock |  |  | Hannum Clock |  |  | Levine Clock |  |  |
| --- | --- | --- | --- | --- | --- | --- | --- | --- | --- | --- | --- | --- |
| | $r$ | 95% CI | p-value | $r$ | 95% CI | p-value | $r$ | 95% CI | p-value | $r$ | 95% CI | p-value |
| Balance | -0.29 | [-0.36, -0.23] | 4.92E-18 | -0.05 | [-0.12, 0.02] | 0.136 | -0.07 | [-0.14, 0.00] | 0.066 | -0.09 | [-0.16, -0.01] | 0.019 |
| Gait Speed | -0.26 | [-0.32, -0.19] | 5.93E-13 | -0.05 | [-0.13, 0.02] | 0.152 | -0.11 | [-0.19, -0.03] | 0.004 | -0.06 | [-0.13, 0.01] | 0.076 |
| Steps in Place | -0.22 | [-0.29, -0.15] | 1.02E-09 | 0.03 | [-0.04, 0.10] | 0.446 | -0.01 | [-0.07, 0.05] | 0.756 | -0.05 | [-0.12, 0.03] | 0.203 |
| Chair Stands | -0.19 | [-0.26, -0.12] | 5.55E-08 | -0.05 | [-0.12, 0.03] | 0.199 | -0.10 | [-0.17, -0.04] | 0.003 | -0.08 | [-0.16, -0.01] | 0.031 |
| Grip Strength | -0.05 | [-0.12, 0.02] | 0.162 | -0.02 | [-0.09, 0.05] | 0.536 | -0.05 | [-0.12, 0.02] | 0.161 | 0.03 | [-0.04, 0.09] | 0.445 |
| Motor Coordination | -0.15 | [-0.21, -0.10] | 6.41E-08 | -0.01 | [-0.06, 0.05] | 0.829 | -0.05 | [-0.11, 0.02] | 0.147 | -0.03 | [-0.09, 0.04] | 0.451 |
| Physical Limitations | 0.20 | [0.11, 0.28] | 1.30E-05 | -0.04 | [-0.11, 0.03] | 0.252 | 0.02 | [-0.04, 0.08] | 0.553 | 0.02 | [-0.04, 0.08] | 0.485 |
| Perceptual Reasoning | -0.28 | [-0.35, -0.22] | 2.38E-17 | -0.02 | [-0.08, 0.05] | 0.611 | -0.13 | [-0.19, -0.07] | 8.25E-05 | -0.05 | [-0.11, 0.02] | 0.162 |
| Working Memory | -0.20 | [-0.27, -0.14] | 8.26E-10 | -0.07 | [-0.14, 0.00] | 0.061 | -0.12 | [-0.19, -0.06] | 3.16E-04 | -0.07 | [-0.13, 0.00] | 0.042 |
| Processing Speed | -0.14 | [-0.21, -0.08] | 1.44E-05 | -0.05 | [-0.12, 0.01] | 0.123 | -0.06 | [-0.13, 0.00] | 0.060 | -0.05 | [-0.11, 0.02] | 0.187 |
| Self-rated Health | -0.27 | [-0.34, -0.20] | 6.83E-13 | -0.04 | [-0.11, 0.04] | 0.327 | -0.05 | [-0.11, 0.02] | 0.172 | -0.13 | [-0.20, -0.06] | 1.98E-04 |
| Facial Aging | 0.35 | [0.28, 0.43] | 3.74E-19 | -0.04 | [-0.11, 0.03] | 0.218 | 0.08 | [0.02, 0.15] | 0.016 | 0.09 | [0.02, 0.16] | 0.009 |
| Adjusted for Estimated Cell Counts at Age 38 |  |  |  |  |  |  |  |  |  |  |  |  |
| Balance | -0.31 | [-0.38, -0.24] | 6.03E-17 | -0.05 | [-0.13, 0.02] | 0.132 | -0.04 | [-0.14, 0.05] | 0.363 | -0.06 | [-0.14, 0.03] | 0.173 |
| Gait Speed | -0.25 | [-0.32, -0.17] | 3.50E-10 | -0.05 | [-0.12, 0.02] | 0.177 | -0.08 | [-0.18, 0.01] | 0.075 | -0.03 | [-0.11, 0.05] | 0.460 |
| Steps in Place | -0.26 | [-0.34, -0.18] | 7.89E-10 | 0.03 | [-0.04, 0.11] | 0.395 | -0.03 | [-0.12, 0.07] | 0.578 | -0.06 | [-0.14, 0.02] | 0.166 |
| Chair Stands | -0.21 | [-0.29, -0.13] | 2.69E-07 | -0.04 | [-0.12, 0.03] | 0.262 | -0.14 | [-0.23, -0.05] | 0.003 | -0.08 | [-0.17, 0.01] | 0.068 |
| Grip Strength | -0.06 | [-0.14, 0.02] | 1.17E-01 | -0.02 | [-0.09, 0.05] | 0.556 | -0.08 | [-0.17, 0.01] | 0.093 | 0.04 | [-0.04, 0.12] | 0.292 |
| Motor Coordination | -0.17 | [-0.22, -0.11] | 6.19E-08 | -0.02 | [-0.07, 0.04] | 0.542 | -0.02 | [-0.10, 0.06] | 0.657 | 0.00 | [-0.07, 0.07] | 0.936 |
| Physical Limitations | 0.21 | [0.12, 0.30] | 8.79E-06 | -0.05 | [-0.12, 0.03] | 0.213 | -0.01 | [-0.10, 0.08] | 0.874 | 0.01 | [-0.06, 0.08] | 0.815 |
| Perceptual Reasoning | -0.28 | [-0.35, -0.21] | 1.60E-14 | -0.02 | [-0.09, 0.05] | 0.565 | -0.09 | [-0.18, 0.00] | 0.045 | 0.01 | [-0.06, 0.09] | 0.748 |
| Working Memory | -0.17 | [-0.24, -0.10] | 5.61E-06 | -0.07 | [-0.14, 0.00] | 0.067 | -0.09 | [-0.19, 0.00] | 0.048 | -0.02 | [-0.10, 0.05] | 0.553 |
| Processing Speed | -0.17 | [-0.24, -0.10] | 4.63E-06 | -0.06 | [-0.13, 0.00] | 0.067 | -0.11 | [-0.20, -0.02] | 0.013 | -0.06 | [-0.14, 0.02] | 0.122 |
| Self-rated Health | -0.29 | [-0.37, -0.21] | 1.99E-12 | -0.05 | [-0.12, 0.02] | 0.199 | -0.03 | [-0.12, 0.06] | 0.475 | -0.14 | [-0.22, -0.06] | 3.09E-04 |
| Facial Aging | 0.36 | [0.28, 0.45] | 5.03E-16 | -0.04 | [-0.11, 0.03] | 0.258 | 0.03 | [-0.06, 0.13] | 0.497 | 0.07 | [-0.01, 0.15] | 0.077 |
| Adjusted for Pack Years at Age 38 |  |  |  |  |  |  |  |  |  |  |  |  |
| Balance | -0.24 | [-0.32, -0.15] | 4.41E-08 | -0.08 | [-0.15, -0.02] | 0.015 | -0.06 | [-0.13, 0.00] | 0.062 | -0.07 | [-0.14, 0.00] | 0.068 |
| Gait Speed | -0.19 | [-0.28, -0.10] | 3.24E-05 | -0.08 | [-0.15, -0.01] | 0.022 | -0.11 | [-0.19, -0.04] | 0.002 | -0.04 | [-0.11, 0.03] | 0.241 |
| Steps in Place | -0.14 | [-0.24, -0.05] | 0.004 | 0.00 | [-0.07, 0.07] | 0.935 | -0.01 | [-0.07, 0.05] | 0.773 | -0.03 | [-0.10, 0.05] | 0.468 |
| Chair Stands | -0.14 | [-0.24, -0.05] | 0.002 | -0.07 | [-0.14, 0.01] | 0.076 | -0.10 | [-0.17, -0.04] | 0.003 | -0.07 | [-0.14, 0.01] | 0.076 |
| Grip Strength | -0.06 | [-0.15, 0.03] | 0.175 | -0.02 | [-0.09, 0.04] | 0.483 | -0.05 | [-0.12, 0.02] | 0.142 | 0.03 | [-0.04, 0.10] | 0.399 |
| Motor Coordination | -0.10 | [-0.17, -0.03] | 0.004 | -0.02 | [-0.08, 0.03] | 0.362 | -0.05 | [-0.11, 0.01] | 0.125 | -0.01 | [-0.08, 0.05] | 0.740 |
| Physical Limitations | 0.15 | [0.05, 0.26] | 0.005 | -0.02 | [-0.09, 0.05] | 0.580 | 0.02 | [-0.04, 0.08] | 0.469 | 0.00 | [-0.06, 0.07] | 0.880 |
| Perceptual Reasoning | -0.21 | [-0.30, -0.13] | 6.17E-07 | -0.05 | [-0.11, 0.02] | 0.136 | -0.13 | [-0.20, -0.07] | 2.58E-05 | -0.02 | [-0.09, 0.04] | 0.469 |
| Working Memory | -0.15 | [-0.23, -0.07] | 3.12E-04 | -0.09 | [-0.16, -0.02] | 0.009 | -0.13 | [-0.19, -0.06] | 2.73E-04 | -0.05 | [-0.11, 0.01] | 0.111 |
| Processing Speed | -0.06 | [-0.14, 0.02] | 0.171 | -0.08 | [-0.14, -0.01] | 0.025 | -0.07 | [-0.13, 0.00] | 0.047 | -0.03 | [-0.09, 0.04] | 0.409 |
| Self-rated Health | -0.18 | [-0.28, -0.09] | 1.20E-04 | -0.07 | [-0.14, 0.00] | 0.054 | -0.05 | [-0.12, 0.02] | 0.132 | -0.10 | [-0.17, -0.04] | 0.002 |
| Facial Aging | 0.24 | [0.14, 0.33] | 1.37E-06 | 0.00 | [-0.07, 0.06] | 0.984 | 0.09 | [0.02, 0.16] | 0.010 | 0.06 | [-0.01, 0.12] | 0.085 |

**Panel B. Analysis of change in healthspan-related characteristics from age 38-45 years**
|  | DunedinPoAm |  |  | Horvath Clock |  |  | Hannum Clock |  |  | Levine Clock |  |  |
| --- | --- | --- | --- | --- | --- | --- | --- | --- | --- | --- | --- | --- |
|  | r | 95% CI | p-value | r | 95% CI | p-value | r | 95% CI | p-value | r | 95% CI | p-value |
| Balance | -0.11 | [-0.19, -0.04] | 2.76E-03 | 0.02 | [-0.06, 0.09] | 0.654 | 0.00 | [-0.08, 0.08] | 0.968 | 0.03 | [-0.04, 0.09] | 0.478 |
| Grip Strength | 0.00 | [-0.07, 0.08] | 0.932 | 0.00 | [-0.07, 0.07] | 0.991 | -0.01 | [-0.09, 0.08] | 0.853 | 0.04 | [-0.03, 0.11] | 0.279 |
| Motor Coordination | -0.05 | [-0.13, 0.02] | 0.147 | -0.01 | [-0.08, 0.06] | 0.802 | -0.03 | [-0.11, 0.06] | 0.539 | -0.04 | [-0.12, 0.04] | 0.381 |
| Physical Limitations | 0.10 | [0.02, 0.18] | 0.016 | -0.07 | [-0.14, 0.01] | 0.102 | -0.03 | [-0.10, 0.03] | 0.335 | -0.05 | [-0.12, 0.02] | 0.129 |
| Cognitive Decline | -0.12 | [-0.19, -0.05] | 7.94E-04 | -0.03 | [-0.10, 0.03] | 0.324 | -0.06 | [-0.14, 0.01] | 0.101 | -0.06 | [-0.13, 0.01] | 0.090 |
| Incident Fair/Poor Health | 1.75 | [1.43, 2.14] | 3.97E-08 | 0.96 | [0.72, 1.27] | 0.758 | 1.07 | [0.84, 1.36] | 0.577 | 1.35 | [1.08, 1.68] | 0.009 |
| Facial Aging | 0.10 | [0.03, 0.18] | 0.005 | -0.04 | [-0.11, 0.03] | 0.279 | -0.01 | [-0.08, 0.06] | 0.762 | 0.05 | [-0.02, 0.11] | 0.146 |
| Adjusted for Estimated Cell Counts at Age 38 |  |  |  |  |  |  |  |  |  |  |  |  |
| Balance | -0.13 | [-0.21, -0.05] | 0.002 | 0.02 | [-0.05, 0.10] | 0.555 | -0.01 | [-0.11, 0.09] | 0.872 | 0.03 | [-0.05, 0.11] | 0.429 |
| Grip Strength | 0.01 | [-0.08, 0.09] | 0.834 | 0.01 | [-0.06, 0.08] | 0.787 | 0.00 | [-0.10, 0.11] | 0.936 | 0.07 | [-0.02, 0.15] | 0.126 |
| Motor Coordination | -0.04 | [-0.12, 0.05] | 0.364 | -0.01 | [-0.08, 0.06] | 0.830 | 0.02 | [-0.09, 0.13] | 0.689 | -0.01 | [-0.09, 0.08] | 0.871 |
| Physical Limitations | 0.10 | [0.01, 0.20] | 0.030 | -0.07 | [-0.15, 0.00] | 0.063 | -0.06 | [-0.15, 0.04] | 0.227 | -0.08 | [-0.16, 0.00] | 0.046 |
| Cognitive Decline | -0.10 | [-0.18, -0.02] | 0.015 | -0.04 | [-0.11, 0.03] | 0.307 | -0.02 | [-0.11, 0.08] | 0.707 | -0.03 | [-0.12, 0.05] | 0.422 |
| Incident Fair/Poor Health | 2.02 | [1.57, 2.59] | 3.03E-08 | 0.97 | [0.73, 1.28] | 0.829 | 1.09 | [0.78, 1.52] | 0.625 | 1.49 | [1.16, 1.92] | 0.002 |
| Facial Aging | 0.09 | [0.01, 0.17] | 0.023 | -0.04 | [-0.11, 0.03] | 0.270 | -0.06 | [-0.15, 0.04] | 0.251 | 0.03 | [-0.05, 0.10] | 0.501 |
| Adjusted for Pack Years at Age 38 |  |  |  |  |  |  |  |  |  |  |  |  |
| Balance | -0.13 | [-0.22, -0.04] | 0.005 | 0.01 | [-0.06, 0.08] | 0.790 | 0.00 | [-0.08, 0.08] | 0.972 | 0.03 | [-0.04, 0.10] | 0.388 |
| Grip Strength | -0.01 | [-0.11, 0.08] | 0.760 | 0.00 | [-0.06, 0.07] | 0.937 | -0.01 | [-0.09, 0.08] | 0.841 | 0.04 | [-0.03, 0.11] | 0.292 |
| Motor Coordination | -0.03 | [-0.12, 0.07] | 0.554 | -0.02 | [-0.09, 0.05] | 0.640 | -0.03 | [-0.11, 0.06] | 0.528 | -0.03 | [-0.11, 0.05] | 0.464 |
| Physical Limitations | 0.10 | [0.00, 0.20] | 0.047 | -0.06 | [-0.14, 0.02] | 0.159 | -0.03 | [-0.09, 0.03] | 0.362 | -0.06 | [-0.13, 0.01] | 0.093 |
| Cognitive Decline | -0.02 | [-0.12, 0.07] | 0.627 | -0.06 | [-0.12, 0.01] | 0.089 | -0.06 | [-0.14, 0.01] | 0.104 | -0.05 | [-0.12, 0.02] | 0.199 |
| Incident Fair/Poor Health | 1.48 | [1.09, 2.00] | 0.011 | 1.05 | [0.80, 1.36] | 0.743 | 1.07 | [0.85, 1.35] | 0.544 | 1.28 | [1.01, 1.61] | 0.040 |
| Facial Aging | 0.09 | [0.00, 0.18] | 0.060 | -0.03 | [-0.10, 0.04] | 0.419 | -0.01 | [-0.08, 0.06] | 0.777 | 0.04 | [-0.02, 0.10] | 0.227 |

**Supplemental Table 4.** Effect-sizes for associations of DunedinPoAm and epigenetic clocks with KDM Biological Age, and self-rated health in the Understanding Society Study. The table reports standardized regression coefficients and their standard errors from linear regression models in which the predictor was the methylation measure listed in the far-left column and the dependent variable was either KDM Biological Age (left side coefficients) or self-rated health (right side coefficients). All models included sex and chronological age as covariates. Model 2 included covariates for cell counts estimated from the methylation data. Model 3 included covariates adjusting for smoking status. Model 4 included nonsmokers only. Prior to analysis, Horvath, Hannum, and Levine Clock values were residualized for chronological age. All methylation variables were residualized for plate number.

|  | KDM Biological Age |  |  | Self-rated Health |  |  |
| --- | --- | --- | --- | --- | --- | --- |
|  | b | 95% CI | p-value | b | 95% CI | p-value |
| **M1. Models adjusted for age and sex** |  |  |  |  |  |  |
| mPoA | 0.200 | [0.15, 0.26] | 1.67E-12 | -0.217 | [-0.27, -0.16] | 1.04E-13 |
| Horvath Clock | 0.050 | [-0.01, 0.11] | 0.090 | -0.003 | [-0.06, 0.05] | 0.921 |
| Hannum Clock | 0.051 | [-0.02, 0.12] | 0.130 | -0.042 | [-0.10, 0.01] | 0.143 |
| Levine Clock | 0.150 | [0.09, 0.21] | 8.26E-07 | -0.088 | [-0.14, -0.03] | 0.001 |
| N |  | 1164 |  |  | 1175 |  |
| **M2. Models adjusted for age, sex, and estimated cell counts** |  |  |  |  |  |  |
| mPoA | 0.186 | [0.13, 0.25] | 2.37E-09 | -0.183 | [-0.25, -0.12] | 1.91E-08 |
| Horvath Clock | 0.061 | [0.00, 0.12] | 0.045 | -0.006 | [-0.06, 0.05] | 0.839 |
| Hannum Clock | 0.034 | [-0.05, 0.12] | 0.425 | -0.006 | [-0.08, 0.07] | 0.867 |
| Levine Clock | 0.151 | [0.09, 0.22] | 5.53E-06 | -0.070 | [-0.13, -0.01] | 0.016 |
| N |  | 1164 |  |  | 1175 |  |
| **M3. Models adjusted for age, sex, and smoking** |  |  |  |  |  |  |
| mPoA | 0.207 | [0.13, 0.28] | 2.78E-08 | -0.155 | [-0.22, -0.09] | 7.44E-06 |
| Horvath Clock | 0.044 | [-0.01, 0.10] | 0.134 | 0.004 | [-0.05, 0.06] | 0.881 |
| Hannum Clock | 0.046 | [-0.02, 0.11] | 0.167 | -0.036 | [-0.09, 0.02] | 0.196 |
| Levine Clock | 0.135 | [0.07, 0.19] | 1.15E-05 | -0.067 | [-0.12, -0.02] | 0.011 |
| N |  | 1160 |  |  | 1171 |  |
| **M4. Models adjusted for age, sex; non-smokers** |  |  |  |  |  |  |
| mPoA | 0.252 | [0.14, 0.36] | 1.05E-05 | -0.141 | [-0.25, -0.03] | 0.014 |
| Horvath Clock | 0.051 | [-0.03, 0.13] | 0.219 | -0.061 | [-0.14, 0.02] | 0.141 |
| Hannum Clock | 0.082 | [-0.01, 0.18] | 0.092 | -0.043 | [-0.13, 0.05] | 0.343 |
| Levine Clock | 0.101 | [0.02, 0.18] | 0.012 | -0.079 | [-0.15, 0.00] | 0.043 |
| N |  | 495 |  |  | 498 |  |

**Supplemental Table 5.** Effect-sizes for associations of DunedinPoAm and epigenetic clocks with morbidity and mortality in the Normative Aging Study. Time-to-event analyses of mortality and chronic disease incidence (Panels A and B) were conducted using Cox proportional hazard models to estimate hazard ratios (HRs) in N=771 participants. Repeated-measures analysis (Panel C) was conducted using Poisson regression within a generalized estimating equations framework to account for the nonindependence of repeated observations of individuals (N=1,488 observations of 771 individuals). All models included covariate adjustment for chronological age. Smoking status was measured from the American Thoracic Society Questionnaire (Ferris, 1978) completed by participants at each assessment wave. We classified participants as being current, former, or never smokers (Gao et al., 2019a). Pack years is the total number of cigarettes smoked across the participants lifetime in units equivalent to the number of cigarettes smoked during a year of smoking 1 pack (20 cigarettes) per day. Age acceleration residuals for epigenetic clocks were calculated by regressing epigenetic age on chronological age and predicting residual values. All methylation measures were standardized to M=0 SD=1 for analysis. Effect-sizes thus reflect risk associated with a 1-SD increase in the methylation measure.

| Panel A. Effect-sizes for time-to-event analysis of mortality |  |  |  |  |  |  |  |  |  |  |  |  |
| --- | --- | --- | --- | --- | --- | --- | --- | --- | --- | --- | --- | --- |
|  | Base Model |  |  | Adjusted for Estimated Leukocyte Distribution |  |  | Adjusted for Smoking Status |  |  | Adjusted for Pack Years |  |  |
|  | HR | 95% CI | p-value | HR | 95% CI | p-value | HR | 95% CI | p-value | HR | 95% CI | p-value |
| DunedinPoAm | 1.29 | [1.16-1.45] | <0.001 | 1.28 | [1.13-1.44] | <0.001 | 1.32 | [1.17-1.48] | <0.001 | 1.26 | [1.12-1.12] | 0.001 |
| Age Acceleration Residuals |  |  |  |  |  |  |  |  |  |  |  |  |
| Horvath Clock | 1.07 | [0.94-1.22] | 0.308 | 1.08 | [0.94-1.24] | 0.284 | 1.07 | [0.94-1.23] | 0.321 | 1.07 | [0.94-1.22] | 0.319 |
| Hannum Clock | 1.05 | [0.90-1.23] | 0.525 | 1.04 | [0.89-1.22] | 0.598 | 1.04 | [0.89-1.22] | 0.599 | 1.02 | [0.87-1.19] | 0.827 |
| Phenotypic Aging Clock | 1.18 | [1.05-1.34] | 0.008 | 1.19 | [1.05-1.35] | 0.008 | 1.18 | [1.04-1.34] | 0.011 | 1.15 | [1.02-1.31] | 0.026 |
| Panel B. Effect-sizes for time-to-event analysis of incident chronic disease |  |  |  |  |  |  |  |  |  |  |  |  |
|  | Base Model |  |  | Adjusted for Estimated Leukocyte Distribution |  |  | Adjusted for Smoking Status |  |  | Adjusted for Pack Years |  |  |
|  | HR | 95% CI | p-value | HR | 95% CI | p-value | HR | 95% CI | p-value | HR | 95% CI | p-value |
| DunedinPoAm | 1.19 | 1.03 - 1.38 | 0.019 | 1.18 | 1.01 - 1.38 | 0.035 | 1.12 | 0.95 - 1.31 | 0.169 | 1.15 | 0.99 - 1.34 | 0.076 |
| Age Acceleration Residuals |  |  |  |  |  |  |  |  |  |  |  |  |
| Horvath Clock | 1.02 | 0.86 - 1.20 | 0.851 | 0.98 | 0.82 - 1.17 | 0.857 | 1.01 | 0.86 - 1.20 | 0.857 | 1.03 | 0.87 - 1.22 | 0.7349 |
| Hannum Clock | 1.17 | 0.99 - 1.39 | 0.066 | 1.16 | 0.97 - 1.39 | 0.094 | 1.18 | 0.99 - 1.40 | 0.062 | 1.17 | 0.99 - 1.39 | 0.0658 |
| Phenotypic Aging Clock | 1.10 | 0.93 - 1.31 | 0.273 | 1.11 | 0.93 - 1.33 | 0.262 | 1.07 | 0.90 - 1.27 | 0.456 | 1.09 | 0.92 - 1.29 | 0.3409 |
| Panel C. Effect-sizes for repeated-measures analysis of prevalent chronic disease |  |  |  |  |  |  |  |  |  |  |  |  |
|  | Base Model |  |  | Adjusted for Estimated Leukocyte Distribution |  |  | Adjusted for Smoking Status |  |  | Adjusted for Pack Years |  |  |
|  | IRR | 95% CI | p-value | IRR | 95% CI | p-value | IRR | 95% CI | p-value | IRR | 95% CI | p-value |
| DunedinPoAm | 1.15 | [1.11-1.20] | <0.001 | 1.15 | [1.10-1.19] | <0.001 | 1.16 | [1.12-1.21] | <0.001 | 1.12 | [1.08-1.17] | <0.001 |
| Age Acceleration Residuals |  |  |  |  |  |  |  |  |  |  |  |  |
| Horvath Clock | 1.07 | [1.03-1.12] | 0.002 | 1.08 | [1.03-1.13] | 0.001 | 1.08 | [1.03-1.13] | 0.001 | 1.07 | [1.03-1.12] | 0.002 |
| Hannum Clock | 1.04 | [1.00-1.09] | 0.074 | 1.03 | [0.98-1.08] | 0.189 | 1.04 | [1.00-1.09] | 0.065 | 1.04 | [0.99-1.09] | 0.090 |
| Phenotypic Aging Clock | 1.13 | [1.08-1.18] | <0.001 | 1.13 | [1.08-1.18] | <0.001 | 1.13 | [1.08-1.18] | <0.001 | 1.12 | [1.07-1.16] | <0.001 |

**Supplemental Table 6.** Effect-sizes for associations of socioeconomic status (SES) and victimization exposure with DunedinPoAm and epigenetic clocks at age 18 in the E-Risk Study. The table shows effect-sizes reported as standardized regression coefficients (b) and 95% confidence intervals (CIs) from models in which childhood family socioeconomic status (SES) and victimization were predictor variables and the dependent variables were the DNA methylation measures. Model 1 included covariate adjustment for sex. Model 2 additionally included covariates for estimated cell counts. (Chronological age was the same for all twins in the birth cohort.) Models 3 and 4 adjusted for smoking status. Only 33% of the analysis sample had ever smoked. Model 3 included covariates adjusting for smoking status at age 18 (never, former, current). Model 4 included nonsmokers only. Methylation measurements were derived from blood DNA methylation collected when Study members were aged 18 years and were residualized for plate number prior to analysis. All E-Risk participants are the same chronological age. Epigenetic clock measures were differenced from this chronological age prior to analysis. Effect-sizes are reported in terms of standard deviation differences in the outcome associated with a 1 standard deviation increase in methylation measures. All models were adjusted for sex. Standard errors were clustered at the family level to account for non-independence of twin data.

|  | DunedinPoAm |  |  | Horvath Clock |  |  | Hannum Clock |  |  | Levine Clock |  |  |
| --- | --- | --- | --- | --- | --- | --- | --- | --- | --- | --- | --- | --- |
|  | b | 95% CI | p-value | b | 95% CI | p-value | b | 95% CI | p-value | b | 95% CI | p-value |
| **M1. Base Model** |  |  |  |  |  |  |  |  |  |  |  |  |
| Childhood SES | -0.18 | [-0.23, -0.13] | 4.70E-11 | -0.01 | [-0.07, 0.04] | 0.633 | 0.01 | [-0.05, 0.07] | 0.720 | -0.08 | [-0.13, -0.02] | 0.007 |
| Victimization | 0.13 | [0.08, 0.18] | 1.42E-06 | 0.00 | [-0.06, 0.05] | 0.924 | -0.02 | [-0.07, 0.04] | 0.577 | 0.01 | [-0.04, 0.07] | 0.617 |
| **M2. Adjusted for estimated cell counts** |  |  |  |  |  |  |  |  |  |  |  |  |
| Childhood SES | -0.17 | [-0.21, -0.12] | 9.45E-14 | -0.02 | [-0.07, 0.03] | 0.428 | 0.01 | [-0.02, 0.04] | 0.559 | -0.07 | [-0.12, -0.03] | 0.002 |
| Victimization | 0.12 | [0.07, 0.16] | 7.30E-07 | 0.00 | [-0.05, 0.05] | 0.900 | -0.02 | [-0.05, 0.01] | 0.236 | 0.01 | [-0.04, 0.05] | 0.760 |
| **M3. Adjusted for smoking status at age 18** |  |  |  |  |  |  |  |  |  |  |  |  |
| Childhood SES | -0.09 | [-0.14, -0.04] | 3.77E-04 | -0.03 | [-0.09, 0.02] | 0.272 | 0.00 | [-0.06, 0.06] | 1.00 | -0.07 | [-0.13, -0.02] | 0.010 |
| Victimization | 0.04 | [-0.01, 0.09] | 0.083 | 0.01 | [-0.04, 0.07] | 0.607 | -0.01 | [-0.06, 0.05] | 0.836 | 0.01 | [-0.05, 0.06] | 0.770 |
| **M4. Non-smokers only** |  |  |  |  |  |  |  |  |  |  |  |  |
| Childhood SES | -0.08 | [-0.13, -0.03] | 0.003 | -0.04 | [-0.10, 0.02] | 0.190 | -0.02 | [-0.08, 0.05] | 0.663 | -0.07 | [-0.14, -0.01] | 0.023 |
| Victimization | 0.07 | [0.01, 0.12] | 0.018 | -0.02 | [-0.09, 0.04] | 0.517 | -0.02 | [-0.09, 0.06] | 0.678 | 0.00 | [-0.07, 0.07] | 0.961 |

**Supplemental Table 7.** Characteristics of participants in the CALERIE Trial and subsample with baseline methylation data. The top panel of the table shows demographic characteristics and measured rate of aging for participants in the Ad Libitum (usual diet) and Caloric Restriction arms of the CALERIE Trial. The middle panel shows the same data for the subset of participants for whom methylation data were available from the baseline CALERIE assessment. The bottom panel shows effect-sizes for associations of methylation measures with the rate of change in KDM Biological Age during follow-up. Effect-sizes are stratified marginal effects computed from regressions of predicted slopes of change in KDM Biological Age on treatment condition, baseline values of the methylation measures, and the interaction of treatment condition and the methylation measures. Effect-sizes for association of baseline DunedinPoAm with rate of change in KDM Biological Age over follow-up are plotted separately by treatment condition (CR for caloric restriction and AL for Ad Libitum control). Effect-sizes reflect the predicted increase in the rate of annual change in KDM Biological Age over the 2 years of follow-up associated with a 1 SD increase in the methylation measure. For example, for DunedinPoaM, a value of 0.2 for participants in the AL control condition indicates that having DunedinPoAm 1 SD higher at baseline is associated with an increase in the aging rate of 0.2 years of physiological change per 12 months of follow-up. Models included covariate adjustment for sex and chronological age at baseline.

**CALERIE Trial**
|  | Ad Libitum (n=75) |  | Caloric Restriction (n=145) |  |
| --- | --- | --- | --- | --- |
|  | M | SD | M | SD |
| Age | 37.86 | 6.94 | 37.89 | 7.32 |
| Male (%) | 0.29 | 0.46 | 0.31 | 0.46 |
| KDM Biological Age | 36.31 | 6.98 | 36.59 | 6.92 |
| Annual Change in KDM Biological Age | 0.72 | 0.95 | 0.11 | 0.96 |

**Sub-sample with baseline methylation data**
|  | Ad Libitum (n=68) |  | Caloric Restriction (n=118) |  |
| --- | --- | --- | --- | --- |
|  | M | SD | M | SD |
| Age | 38.16 | 7.07 | 38.37 | 7.30 |
| Male (%) | 0.31 | 0.47 | 0.31 | 0.47 |
| KDM Biological Age | 36.46 | 6.99 | 37.19 | 6.73 |
| Annual Change in KDM Biological Age | 0.73 | 0.95 | 0.12 | 1.03 |
| mPoA | 1.00 | 0.05 | 1.01 | 0.06 |

|  | DunedinPoAm |  |  | Horvath Clock |  |  | Hannum Clock |  |  | Levine Clock |  |  |
| --- | --- | --- | --- | --- | --- | --- | --- | --- | --- | --- | --- | --- |
|  | b | 95% CI | p-value | b | 95% CI | p-value | b | 95% CI | p-value | b | 95% CI | p-value |
| AL Control Group | 0.22 | [-0.05, 0.49] | 0.105 | -0.23 | [-0.45, 0.00] | 0.049 | -0.16 | [-0.45, 0.13] | 0.283 | -0.03 | [-0.22, 0.17] | 0.772 |
| CR Intervention Group | -0.08 | [-0.24, 0.09] | 0.351 | 0.07 | [-0.12, 0.27] | 0.443 | -0.11 | [-0.28, 0.05] | 0.183 | 0.02 | [-0.16, 0.21] | 0.796 |
| **Adjusted for estimated cell counts** |  |  |  |  |  |  |  |  |  |  |  |  |
| AL Control Group | 0.23 | [-0.05, 0.52] | 0.105 | -0.22 | [-0.47, 0.02] | 0.074 | -0.22 | [-0.58, 0.14] | 0.225 | 0.01 | [-0.22, 0.25] | 0.919 |
| CR Intervention Group | -0.08 | [-0.27, 0.11] | 0.424 | 0.09 | [-0.11, 0.29] | 0.381 | -0.20 | [-0.47, 0.07] | 0.154 | 0.08 | [-0.15, 0.30] | 0.496 |

**Supplemental Figure 1.**
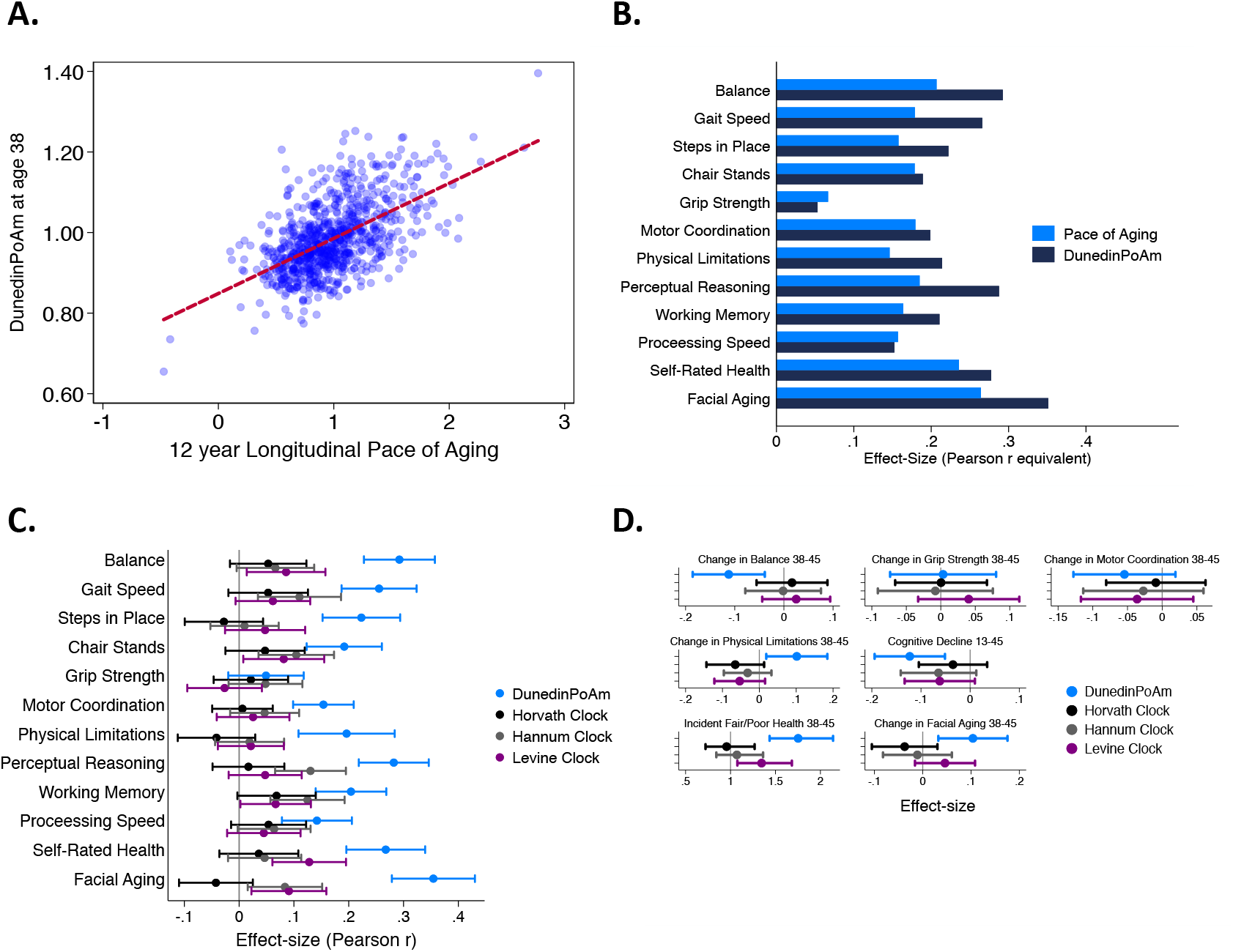
Association of DunedinPoAm with Pace of Aging in the Dunedin Study and Comparison of Effect-sizes for DunedinPoAm, Pace of Aging, and epigenetic clocks. Panel A shows a scatterplot and fitted regression slope for the association between DunedinPoAm trained in the full Dunedin sample and the Pace of Aging measure that was used as the criterion in the training analysis. The correlation of r=0.56 is a within-training sample estimate of association and therefore reflects the upper-bound of possible true association between DunedinPoAm and Pace of Aging. The figure shows data for the N=810 Dunedin Study members included in the training analysis. Panel B graphs effect-sizes for associations of age-38 DunedinPoAm and 12-year longitudinal Pace of Aging with measures of physical and cognitive functioning and subjective signs of aging measured at the Dunedin Study age-45 assessment. Panels C and D plot effect-sizes for DunedinPoAm and the epigenetic clocks proposed by Horvath, Hannum, and Levine for analysis of age-45 outcomes (Panel C) and change from age 38 to 45 (Panel D). Effect-sizes are standardized regression coefficients (interpretable as Pearson r). Effect-sizes are computed for “epigenetic-age acceleration” values of the clocks (i.e. epigenetic ages residualized for chronological age). Error bars show 95% CIs. For all outcomes, effect-sizes are largest for DunedinPoAm.

**Supplemental Figure 2.**
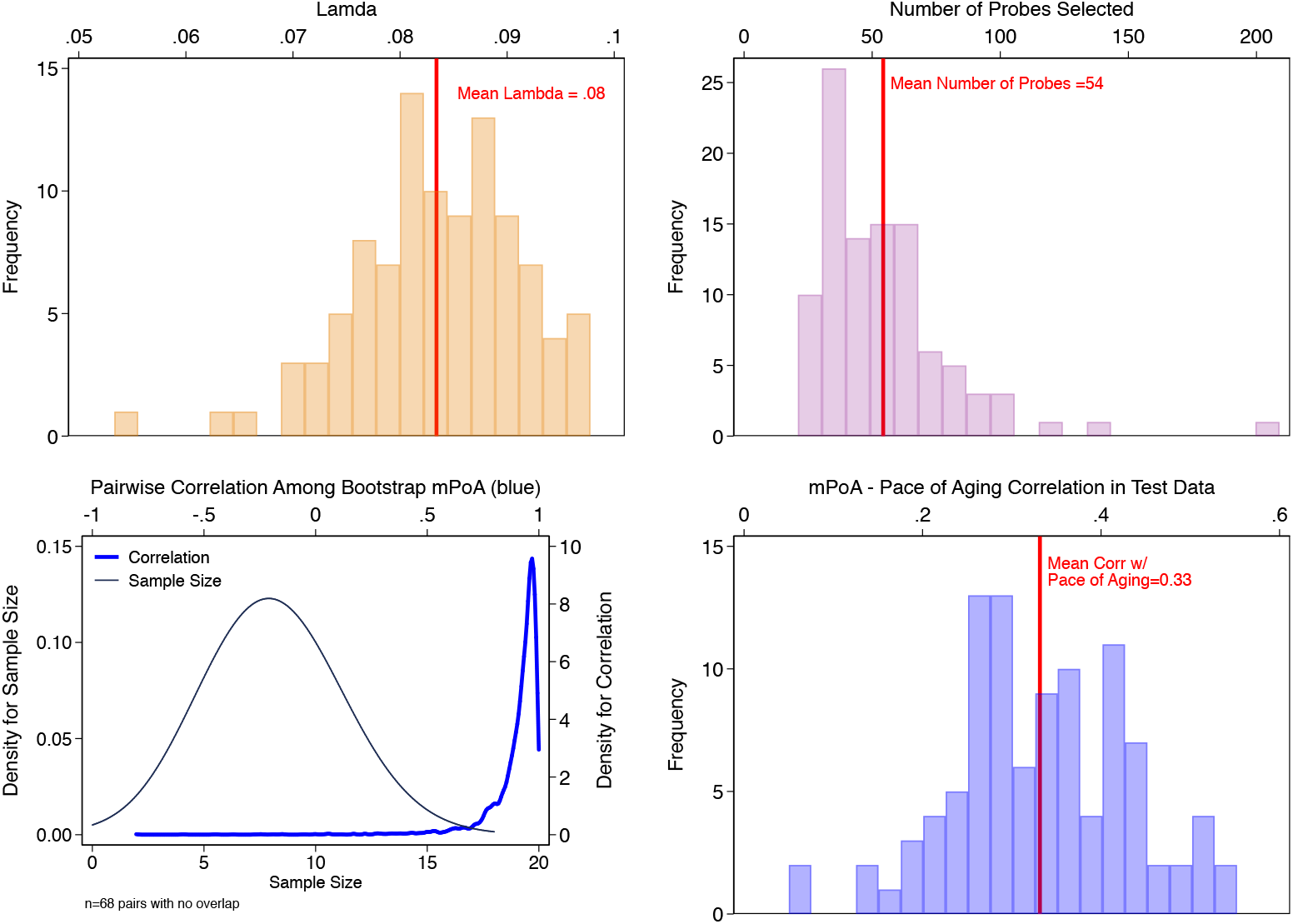
Bootstrap repetition analysis to estimate out-of-sample correlation between mPoA and longitudinal Pace of Aging. The figure shows results from our Bootstrap crossfold validation analysis to evaluate mPoA within the Dunedin Study. Panel A (top left) shows the distribution of elastic-net regression Lambda values estimated across the 100 bootstrap training samples. Panel B (top right) shows the distribution of the number of probes selected by the elastic net regression to compose mPoA across the 100 bootstrap training samples. Panel C (bottom left) graphs two densities illustrating analysis of intercorrelations among the different mPoA algorithms estimated across the bootstrap repetitions. The first density (thin gray line, left side Y axis) shows the distribution of sample sizes for calculation of correlations between pairs of mPoA algorithms. There were 68 pairs of mPoA algorithms for which there were no overlapping samples (i.e. sample size=0). The second density (thick blue line, right side Y axis) shows the distribution of pairwise correlations. Nearly all of the correlations were r>0.95. Panel D (bottom right) shows the distribution of correlations between mPoA and Pace of Aging in the 100 bootstrap test samples.

**Supplemental Figure 3.**
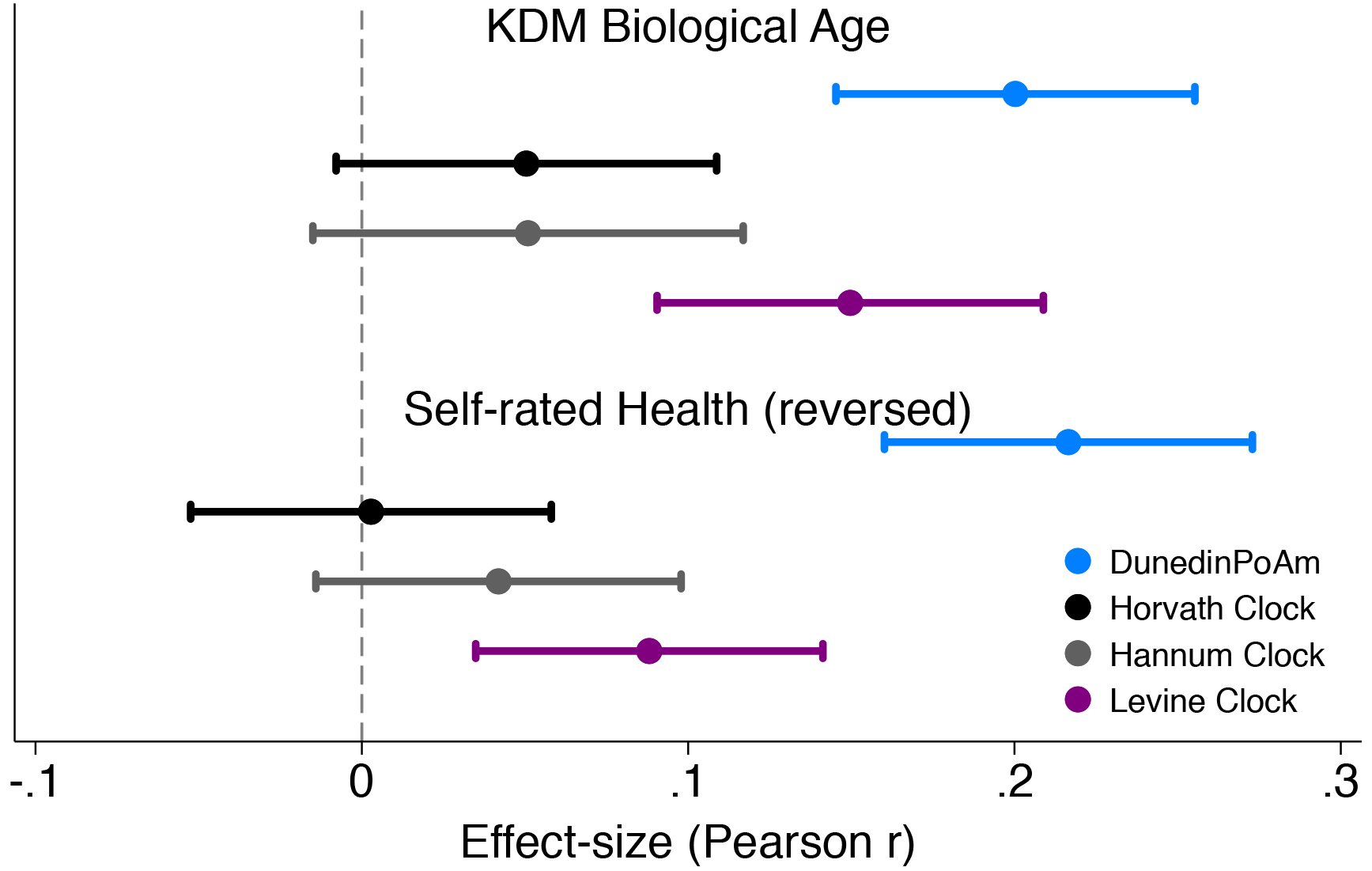
Effect-sizes for associations of DunedinPoAm and epigenetic clocks with KDM Biological Age and self-rated health in Understanding Society. Effect-sizes are standardized regression coefficients (interpretable as Pearson r). Effect-sizes are computed for “epigenetic-age acceleration” values of the clocks (i.e. epigenetic ages residualized for chronological age). The dependent variable in analysis of KDM Biological Age was the difference between KDM Biological Age and chronological age. Self-rated health scores were reversed for analysis so that higher values correspond to ratings of poorer health. All models were adjusted for chronological age and sex.

**Supplemental Figure 4.**
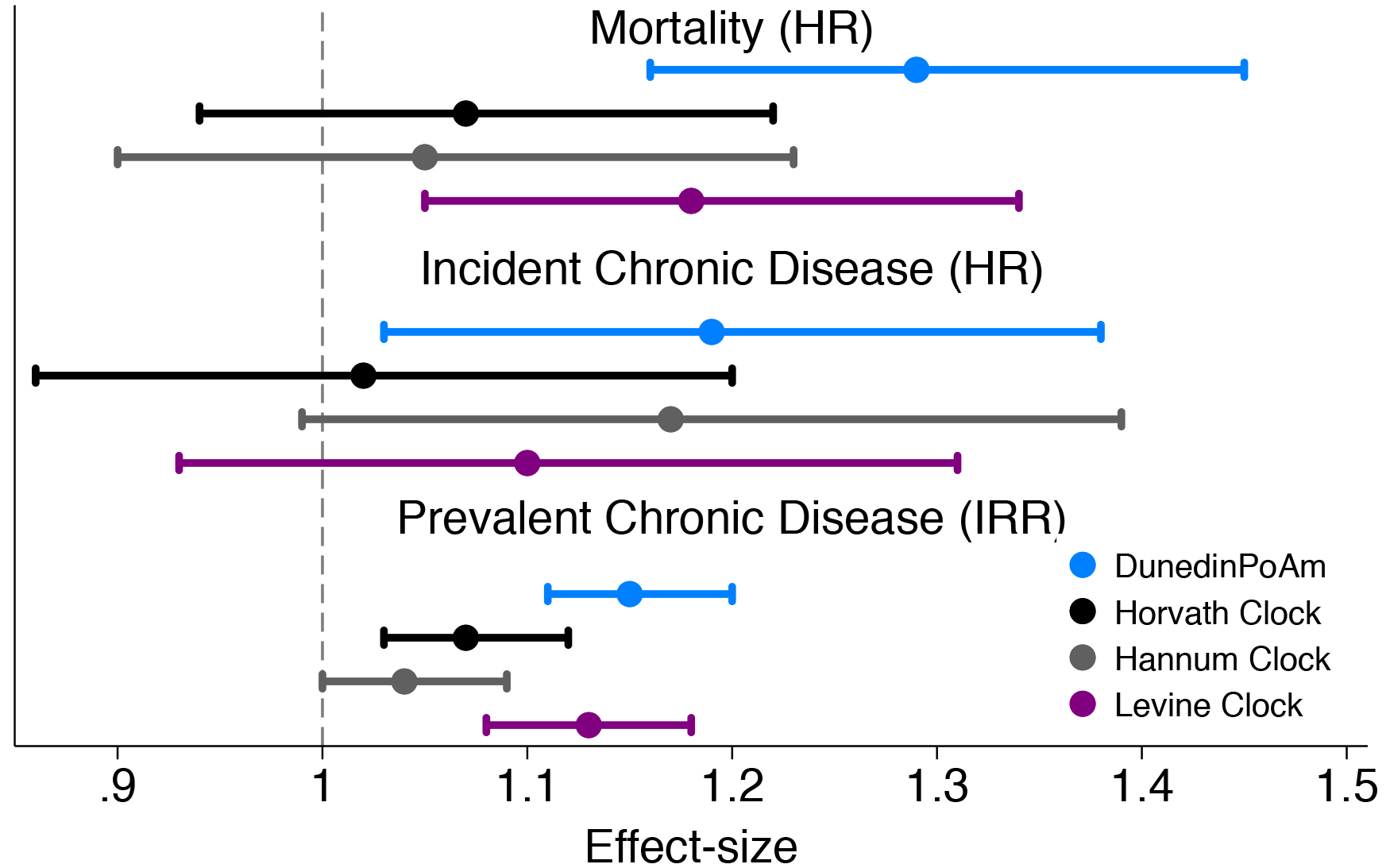
Effect-sizes for association of DunedinPoAm and epigenetic clocks with mortality and morbidity in the Normative Aging Study. Time-to-event analysis of mortality and chronic disease incidence were conducted using Cox proportional hazard models to estimate hazard ratios (HRs) in N=771 participants. Repeated-measures analysis of chronic disease prevalence was conducted using Poisson regression to estimate incidence rate ratios (IRRs) within a generalized estimating equations framework to account for the nonindependence of repeated observations of individuals (N=1,488 observations of 771 individuals). All models included covariate adjustment for chronological age. Age acceleration residuals for epigenetic clocks were calculated by regressing epigenetic age on chronological age and predicting residual values. All methylation measures were standardized to M=0 SD=1 for analysis. Effect-sizes thus reflect risk associated with a 1-SD increase in the methylation measure.

**Supplemental Figure 5.**
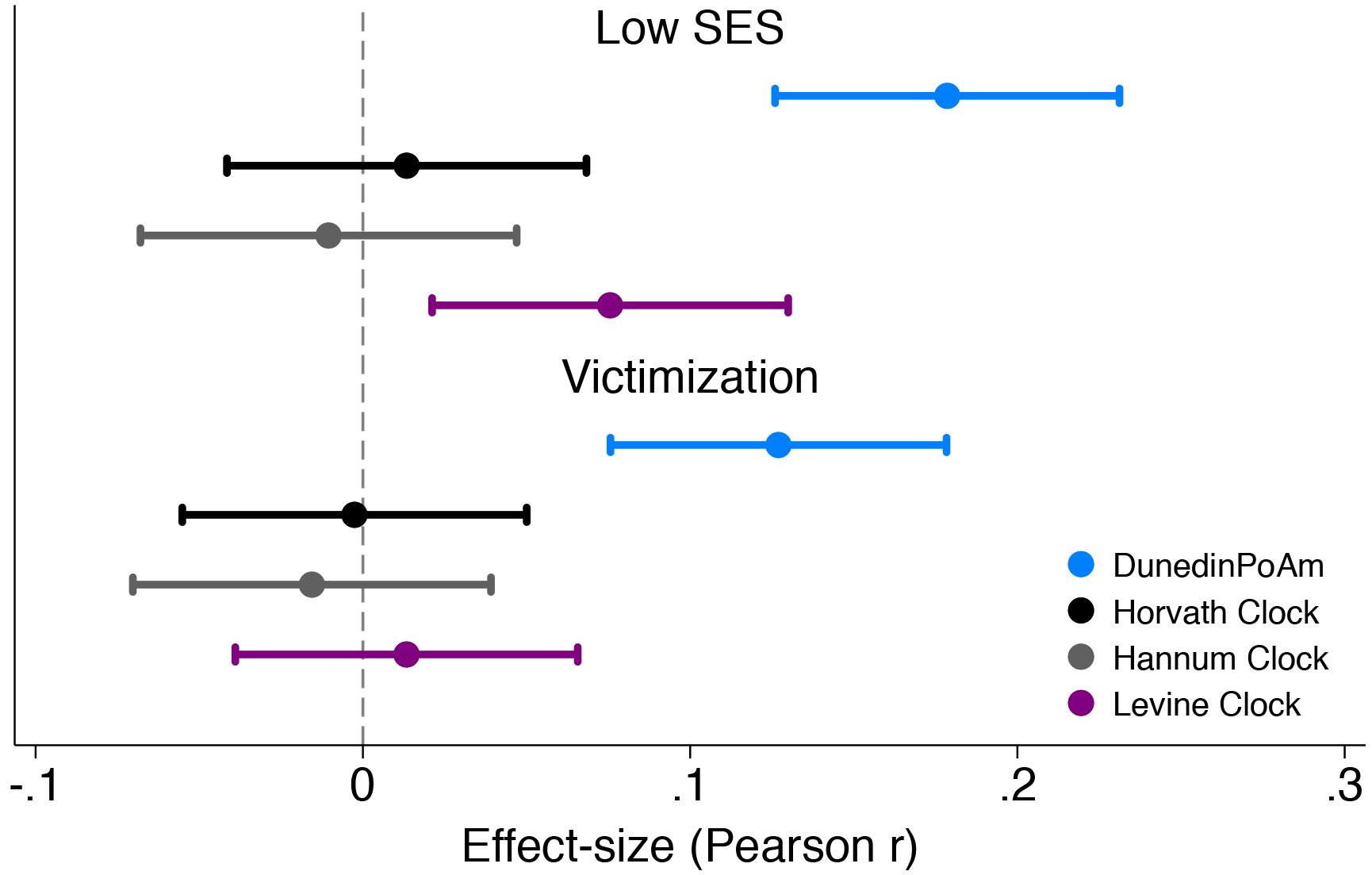
Effect-sizes for associations of childhood socioeconomic status and victimization exposure with DunedinPoAm and epigenetic clocks. Effect-sizes are standardized regression coefficients and 95% confidence intervals from models in which childhood family socioeconomic status (SES) and victimization were predictor variables and the dependent variables were the methylation measures. All E-Risk participants were the same chronological age (18 years) at the time of blood collection. Epigenetic clock measures were differenced from this chronological age prior to analysis. Effect-sizes are reported in terms of standard deviation differences in the methylation measures associated with a 1 standard deviation increase in the predictor measure. All models were adjusted for sex. Standard errors were clustered at the family level to account for non-independence of twin data.

**Supplemental Figure 6.**
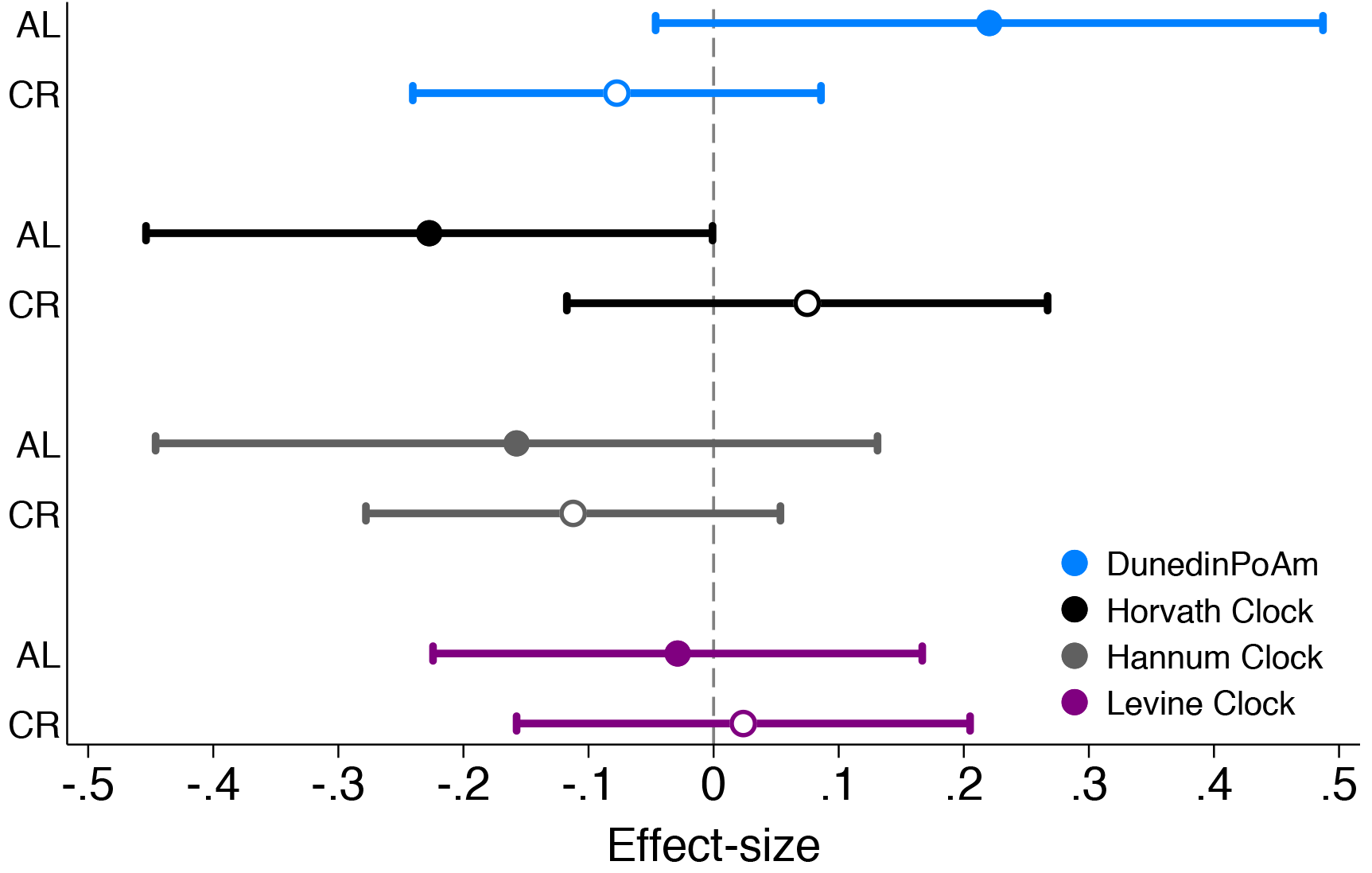
Effect-sizes for associations of DunedinPoAm and epigenetic clocks with future rate of change in KDM Biological Age in Ad Libitum (AL) control group and Caloric Restriction (CR) intervention group participants in the CALERIE Trial. Effect-sizes are stratified marginal effects computed from regressions of predicted slopes of change in KDM Biological Age on treatment condition, baseline values of the methylation measures, and the interaction of treatment condition and the methylation measures. Effect-sizes for association of baseline DunedinPoAm with rate of change in KDM Biological Age over follow-up are plotted separately by treatment condition (AL for Ad Libitum control, plotted as solid-colored circles, CR for caloric restriction, plotted as hollow circles). Effect-sizes reflect the predicted increase in the rate of annual change in KDM Biological Age over the 2 years of follow-up associated with a 1 SD increase in the methylation measure. For example, for DunedinPoaM, a value of 0.2 for participants in the AL control condition indicates that having DunedinPoAm 1 SD higher at baseline is associated with an increase in the aging rate of 0.2 years of physiological change per 12 months of follow-up. Models included covariate adjustment for sex and chronological age at baseline.

